# Lymphotoxin-driven cancer cell eradication by tumoricidal CD8^+^ TIL

**DOI:** 10.1101/2025.11.19.689204

**Authors:** Hongyan Xie, Aiping Jiang, Aonkon Dey, Joseph W. Dean, Jonathan J. Perera, Neal P. Smith, Alex C. Y. Chen, Seth Anderson, Angelina M. Cicerchia, Yi Sun, William A. Michaud, Maria Florentin, Jacy Fang, Or-Yam Revach, Tatyana Sharova, Aleigha Lawless, Katherine H. Xu, Yuhui Song, Bidish Patel, Jonathan D. Stevens, William J. Lane, Derin B. Keskin, Sonia Cohen, Donald P. Lawrence, Ryan J. Sullivan, Keith T. Flaherty, Genevieve M. Boland, Linda T. Nieman, Moshe Sade-Feldman, Nir Hacohen, Debattama R. Sen, Catherine Wu, Brian Gastman, Rongsu Qi, Hequn Yin, Alexandra-Chloé Villani, Robert T. Manguso, Russell W. Jenkins

**Affiliations:** Mass General Cancer Center, Krantz Family Center for Cancer Research, Department of Medicine, Massachusetts General Hospital, Boston, MA, USA; Harvard Medical School, Boston, MA, USA; Broad Institute of MIT and Harvard, Cambridge, MA, USA; Center for Immunology and Inflammatory Diseases, Massachusetts General Hospital, Boston, MA, USA; Iovance Biotherapeutics, Inc., San Carlos, CA, USA; Division of Gastrointestinal and Oncologic Surgery, Department of Surgery, Massachusetts General Hospital, Boston, MA, USA; Department of Pathology, Brigham and Women’s Hospital, Boston, MA 02215, USA; Department of Medical Oncology, Dana-Farber Cancer Institute, Boston, MA, USA; Translational Immunogenomics Laboratory, Dana-Farber Cancer Institute, Boston, Massachusetts, USA; Department of Computer Science, Metropolitan College, Boston University, Boston, Massachusetts, USA; Section for Bioinformatics, Department of Health Technology, Technical University of Denmark, Lyngby, Denmark; Mass General Cancer Center, Department of Medicine, Massachusetts General Hospital, Boston, MA, USA; Hollings Cancer Center, Medical University of South Carolina, Charleston, South Carolina, USA

**Keywords:** tumor-infiltrating lymphocytes, melanoma, lymphotoxin, interferon, cell death

## Abstract

Tumor-infiltrating lymphocyte (TIL) therapy is FDA-approved for patients with treatment-resistant advanced melanoma, but the TIL subpopulations critical for tumor eradication remains incompletely understood. Using patient-derived TIL-melanoma co-cultures, we identified and characterized a novel subset of CD8^+^ TIL, capable of class I HLA-independent cancer cell lysis. The lymphotoxin β receptor (LTβR) and interferon (IFN) sensing pathways were nominated as key determinants of TIL-mediated cancer cell killing from a whole-genome, loss-of-function CRISPR screen. Validation studies confirmed that dual LTβR and IFN sensing is necessary and sufficient for cancer cell lysis, and that expanded CD8^+^ TIL express high lymphotoxin β (*LTB*) and upregulate lymphotoxin α (*LTA*) upon coculture with cancer cells. Leveraging paired scRNA-seq and scTCR-seq data, we confirmed that enrichment of *LTB*^+^*CD8*^+^ T cells is associated with clinical response to TIL, and that *LTB*^+^*CD8*^+^ TIL are expanded from putative neoantigen-reactive, *LTB*^lo^ CD8^+^ T cells in resected tumors.

**Significance:** We have uncovered a previously unrecognized mechanism of TIL-mediated tumor eradication, providing mechanistic insights into the role of LTBR/IFN signaling in TIL-mediated cancer cell killing, and potentially offering insights into novel strategies to isolate, enrich, and expand tumoricidal TIL or augment specific TIL functions to enhance tumor control.

## Introduction

Despite the success of immune checkpoint blockade (ICB) cancer immunotherapy in melanoma and other cancers, resistance to ICB remains common (1,2). Adoptive cell therapy (ACT) using autologous tumor-infiltrating lymphocytes (TIL) was FDA approved in early 2024 and is now available for patients with ICB-resistant melanoma. ACT with TIL involves surgical resection of a metastatic lesion, *in vitro* expansion of autologous TIL using specific growth factors (e.g., interleukin-2) over several weeks, and ultimately reinfusion of billions of autologous TIL back into the patient (3). Objective response rates to TIL therapy of 30-50% have been observed in heavily pre-treated melanoma patients for whom therapeutic options are otherwise limited (4).

While advances in single-cell RNA-sequencing (scRNA-seq) have identified key T cell states associated with response to ICB and ACT (5,6), the precise identity of the “tumoricidal” sub-population of T cells responsible for recognition and elimination of cancer cells and durable clinical response has remained unclear. CD8^+^ T cell states associated with clinical benefit to ICB and ACT are defined by increased expression of genes associated with stem/memory-like T cells (e.g., *TCF7*), with relatively lower expression of genes and gene programs involved in effector T cell function and are relatively diverse with low TCR clonality (5,6). On the other hand, most “tumor-reactive” and clonally expanded CD8^+^ T cells express high levels of dysfunction/exhaustion markers and are hypoproliferative with diminished effector function. Given that multiple studies have demonstrated that T cell exhaustion is associated with irreversible epigenetic changes (7–9), it remains unknown whether clonally expanded, terminally exhausted CD8^+^ T cells directly contribute to anti-tumor immune responses.

Cytotoxic CD8^+^ T cells classically eliminate virus-infected or cancer cells via recognition of an antigenic peptide presented by class I major histocompatibility complex (pMHC) molecules on cancer cells by the T cell receptor (TCR) (22). Low or absent expression of the beta-2 microglobulin (*B2M*) subunit of MHC class I (MHC-I) with resultant impaired surface expression of class I MHC is a reported ICB resistance mechanism (23,24). Unexpectedly, patients with diminished expression of *B2M* also showed ICB responsiveness (25–27), suggesting the existence of MHC-I independent mechanisms of tumor control.

Lymphotoxin β receptor (LTβR), a member of the TNFR superfamily, is expressed on various immune and cancer cells but not on lymphocytes, and binds heterotrimers of LTα/LTβ (LTα1β2) as well as LIGHT (TNFSF14, CD258, HVEM-L) (10–12). LTβ, a type II membrane protein, forms heterotrimers with soluble LTα (*LTA*; formerly known as TNFβ) to engage LTβR (10,13). Activation of LTβR signaling has been implicated in driving chemokine production (14,15), lymphoid tissue development (16,17), and shaping of the tumor microenvironment (18). Additionally, LTβR engagement on tumor cells, particularly with IFNγ, can directly trigger cancer cell death (19–21), yet the contribution of lymphotoxin-expressing CD8⁺ T cells to this process remains unclear.

Here, we demonstrate that a tumoricidal subpopulation of TIL expressing *LTB* can eliminate cancer cells in an MHC-I-independent fashion via dual activation of LTβR- and JAK-STAT pathways in cancer cells. Tumoricidal *LTB*^+^ CD8^+^ TIL arise from tumor-infiltrating CD8^+^ T cells with low or absent *LTB* expression, are enriched after *in vitro* T cell expansion, and are associated with clinical response to TIL therapy.

### Patient-derived TIL-melanoma co-cultures

To examine the cytolytic capacity of TIL using clinically relevant specimens, we generated a cohort of paired TIL and cancer cells (*n* = 10) from patients with advanced melanoma, nine of whom had ICB-resistant disease (**Fig. 1A-B, Methods**). Melanoma cell lines were validated by sequencing which confirmed stereotypical melanoma mutations (e.g., *BRAF*, *NRAS*, *TERT*) (**Fig. 1B and Supplementary Table S1**). The expanded TIL consisted of predominantly CD8^+^ T cells or a mixed population of CD4^+^ and CD8^+^ T cells **(Fig. 1C and Supplementary Fig. S1A-K**, as previously described (28)). Examination of co-inhibitory receptor expression profiles on CD8^+^ T cells demonstrated variable patterns of expression of CD39, TIM-3, and PD-1 (**Fig. 1C and Supplementary Fig. S1A-K**).

**Figure 1.**
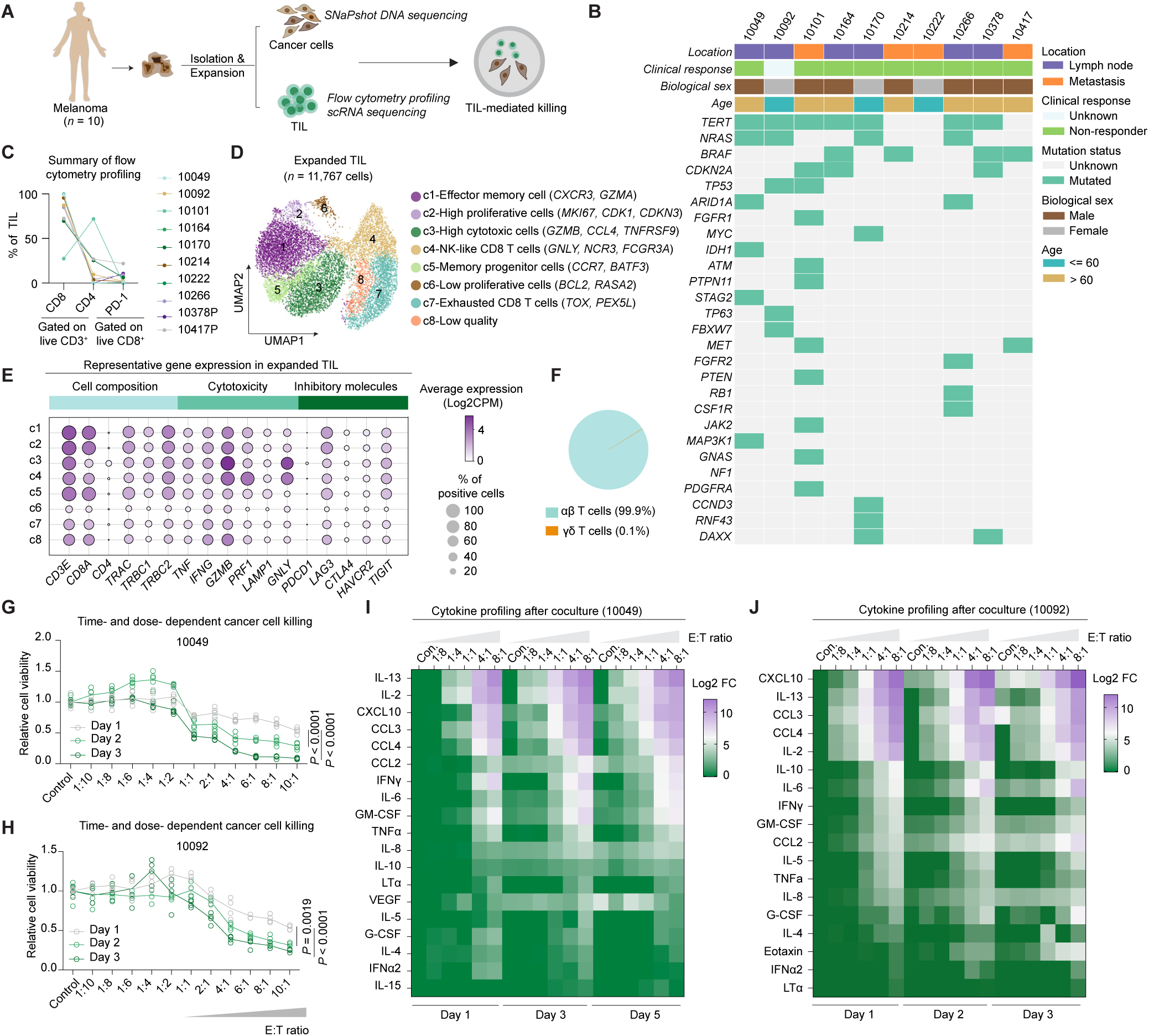
| Patient-derived TIL-melanoma co-cultures. **A**, Schematic of workflow to establish patient-derived melanoma and TIL cocultures. **B**, CoMut plot showing the location, mutation status, clinical response, biological sex, and age of each patient included in this study. **C,** Summary of the immunophenotyping (CD4, CD8, PD-1) of expanded TIL. **D**, Uniform manifold approximation and projection (UMAP) projection of expanded CD8^+^ TIL (*n* = 11,767 cells) with 8 unique populations identified from scRNA-seq analysis of *n* = 6 TIL (10049, 10101, 10214, 10378, 10378P, 10417P). **E**, Dot plot for selected genes expressed in the eight subclusters of TIL. **F**, Percentage of ɑβ and γδ T cells in expanded TIL inferred from scRNA-seq analysis. **G-H**, Viability assessment of (**G**) 10049 and (**H**) 10092 melanoma cells cocultured with autologous TIL at the indicated effector:target (E:T) ratios and time points (*n* = 4-5; Two-way ANOVA with Tukey correction of multiple comparisons). **I-J**, Heatmaps of secreted cytokine profiles of conditioned media after coculture of (**I**) 10049 and (**J**) 10092 melanoma cells and their autologous CD8^+^ TIL at the indicated E:T ratios and time points. Log2 fold change (Log2FC) normalized to control groups is color-coded in the heatmaps.

Next, we performed scRNA-seq (*n* = 11,767 cells) to deeply characterize the subpopulation(s) of expanded TIL and identified eight unique CD8^+^ T cell subpopulations after unbiased clustering (29) (**Fig. 1D, Supplementary Fig. S2A-B**). Clusters 1-5 (effector memory, proliferative, cytotoxic, NK-like, memory progenitor) of expanded TIL were notable for increased expression of genes involved in T cell cytotoxicity and effector function, including *TNF*, *IFNG*, *GZMB*, *PRF1*, and *LAMP1* (encoding CD107a) (**Fig. 1D-E**). Clusters 6-8 were composed of TIL with features of low proliferative capacity (c6), features of exhaustion (c7), and low quality (c8). Expression of genes that encode immune inhibitory molecules, such as *PDCD1* (encoding PD-1), *HAVCR2* (encoding TIM3), and *CTLA4* was expressed at low levels in all clusters, with moderate expression of *LAG3* and *TIGIT* (**Fig. 1E)**, consistent with our flow cytometry analysis **(Fig. 1C, Supplementary Fig. S1A-K**). TCR gene analysis demonstrated that the expanded TIL are largely conventional aβ CD8^+^ T cells expressing *TRAC, TRBC1,* and *TRBC2* (**Fig. 1E-F**) with a low percentage of unconventional CD8^+^ T cells, such as CD1d-restricted nature killer T (NKT) cells, mucosal-associated invariant T (MAIT) cells, and γδ T cells, as evidenced by the low or absent expression of *TRAV1-2* (encoding the invariant TCR chain for MAIT cells (30)), *TRDC*, *TRDV1*, and *TRDV2* (encoding the invariant TCR chains for γδ T cells (25)) (**Fig. 1E-F, Supplementary Fig. S2C**).

To determine the cytotoxic capacity of TIL towards matched melanoma cells, we examined the viability of cancer cells after co-culture with autologous TIL from two patients (10049, 10092) using different effector:target (E:T) ratios and observed time- and dose-dependent cancer cell killing (**Fig. 1G-H**). Bead-based cytokine profiling of conditioned medium from TIL-melanoma cell cocultures confirmed time- and dose-dependent elaboration of multiple cytokines (e.g., IL-2, IL-13) and chemokines (e.g., CCL3, CCL4, CXCL10) as well as effector cytokines (e.g., IFNγ, TNFα, LTα) (**Fig. 1I-J)**. These data demonstrated that expanded TIL from ICB-resistant melanoma patients are capable of eliminating autologous cancer cells.

### MHC I-independent killing of cancer cells by TIL

Given the essential role of pMHC-TCR interaction in T cell activation and recognition (31,32), we next sought to confirm the requirement for MHC-I presentation in TIL-mediated cancer cell lysis. We first examined the effect of pretreating melanoma cells with anti-MHC-I neutralizing antibodies, which effectively blocked the recognition of a pan-HLA-class I antibody detected by flow cytometry (**Fig. 2A**), but surprisingly failed to prevent tumor cell lysis following TIL treatment in 8 TIL-cancer cell pairs (**Fig. 2B**). To confirm this observation, we performed CRISPR/Cas9-gene editing to delete *B2M* as an alternate means of reducing class I MHC surface expression to disrupt the pMHC-TCR interaction. Consistent with our initial observations, loss of *B2M* did not impact TIL-mediated cancer cell killing, as each pair (*n* = 8) of control sgRNA and *B2M* sgRNA melanoma cells exhibited similar sensitivity to TIL (**Fig. 2C, top**) despite near complete loss of surface MHC-I expression (**Fig. 2C, bottom**). Time- and dose-dependent lysis of control and *B2M*-null melanoma cells was observed following co-culture with matched TIL (**Supplementary Fig. S2D-K**). Interestingly, preferential killing of control cancer cells was observed at a lower E:T ratio (1:1) in 2 of 7 pairs, whereas comparable cancer cell killing was observed in all 7 examined TIL-cancer cell pairs at higher E:T ratios (2:1 or 4:1) and longer co-culture time (3 days) (**Supplementary Fig. S2D-K**).

**Figure 2.**
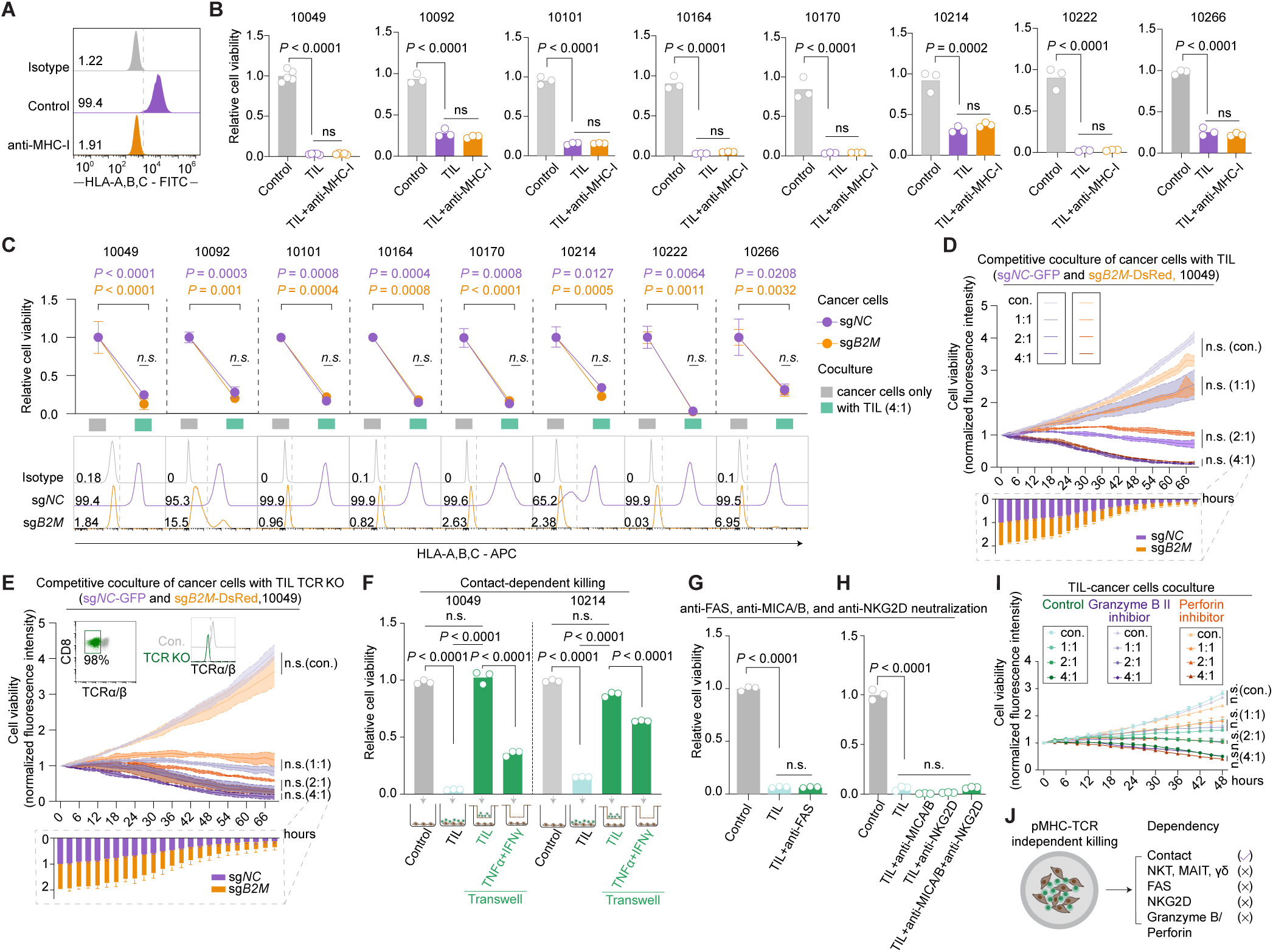
| MHC I-independent killing of cancer cells by TIL. **A**, Expression of HLA-A, B, C in 10049 melanoma cells with/without the pretreatment of MHC-I neutralizing antibodies (20 μg/mL, 3 hours). **B,** Cell viability of 10049, 10092, 10101, 10164, 10170, 10214, 10222, and 10266 melanoma cells with/without the pretreatment of MHC-I neutralizing antibodies cocultured with autologous TIL at the E:T ratio of 0:1 or 4:1 for 2 days. Individual values (open circles) are shown (*n* = 3; one-way ANOVA with Tukey correction of multiple comparisons). **C**, *Top*: viability of control (sg*NC*) and *B2M*-null (sg*B2M*) melanoma cell lines (*n* = 8) cocultured with autologous TIL at E:T ratios of 0:1 or 4:1 for 2 days. Mean +/− s.d. (bar) was shown (*n* = 3-4, 2-sided paired *t-*test). *Bottom*: HLA-A, B, C expression by flow cytometry in control (sg*NC*) and *B2M*-null (sg*B2M*) melanoma cells. **D-E**, Competition assay using equal proportions of control (GFP) and *B2M*-null (DsRed) 10049 melanoma cells co-cultured with (**D**) autologous TIL or (**E**) TCR KO (sg*TRAC* and sg*TRBC*) autologous TIL at different E:T ratios over indicated time points. Normalized fluorescence intensity of viable cancer cells (mean +/− s.d.; shaded region) is shown (*n* = 3, two-way ANOVA with Tukey correction of multiple comparisons; *n.s.*, not significant). Inset stacked bar blot (E:T of 4:1) depicts proportional decrease in control (GFP) and *B2M*-null (DsRed) 10049 melanoma cells over time (*n* = 3, two-way ANOVA with Tukey correction of multiple comparisons). **F**, Viability assessment of indicated melanoma cell lines cocultured with autologous TIL (E:T 4:1) or 40 ng/mL IFNγ plus 160 ng/mL TNFα on either side of a 0.4 μm transwell for 2 days compared to untreated control. Individual values (open circles) are shown (*n* = 3; one-way ANOVA with Tukey’s multiple-comparison test; *n.s.*, not significant). **G-H**, Viability assessment of melanoma cells (10049) cocultured with autologous TIL pretreated with (**G**) anti-FAS or (**H**) anti-MICA/B and/or anti-NKG2D neutralizing antibodies and then cocultured with TIL (E:T 4:1) for 2 days compared to untreated control. Individual values (open circles) are shown (*n* = 3; one-way ANOVA with Tukey correction of multiple comparisons, *n.s.*, not significant.). **I**, Normalized fluorescence intensity of viable 10049-GFP melanoma cells +/−treatment with the Granzyme B II inhibitor (15 nM) or perforin inhibitor (Perforin_IN-2, 10 µM) cocultured with autologous TIL at the indicated E:T ratios for 2 days (*n* = 3; Two-way ANOVA with Tukey correction of multiple comparisons). **J**, Summary of dependency for pMHC-TCR independent killing in TIL-cancer cocultures.

To further evaluate these findings, we performed a time-lapse, image-based competition assay using equal proportions of control (GFP) and *B2M*-null (DsRed) melanoma cells co-cultured with autologous TIL at different E:T ratios over several days. Consistent with previous results, no significant difference in sensitivity to TIL treatment was observed between control (GFP) and *B2M*-null (DsRed) melanoma cells (10049, 10378), especially at higher E:T ratios (2:1 or 4:1) and longer co-culture times (**Fig. 2D, Supplementary Fig. S2L-N**). Moreover, TIL lacking TCRα/β (sg*TRAC* and sg*TRBC*) also exhibited comparable activity against control (GFP) and *B2M*-null (DsRed) melanoma cells (10049, 10378) compared to control TIL (**Fig. 2E**, **Supplementary Fig. S2L-O**). Taken together, these findings demonstrated that TIL can eliminate cancer cells independent of pMHC-TCR interactions.

### TIL-mediated killing requires cell contact, but not unconventional T cells, FAS-FASL, NKG2D-NKG2DL, or granzyme B/perforin

As TIL secrete effector cytokines capable of inducing cancer cell death (e.g., TNFα, IFNγ), we next examined whether lysis of cancer cells required physical contact with TIL. The efficacy of autologous TIL (10049, 10214) was completely abolished when TIL and melanoma cells were placed on opposite sides of a transwell system that was restrictive to passage of TIL, but permissive to smaller soluble molecules (e.g., cytokines) (**Fig. 2F**). While recombinant cytokines (TNFɑ/IFNγ) effectively passed through the transwell and induced cancer cell death (**Fig. 2F**), we observed no effect of conditioned media collected 48 hours after co-culture of TIL with matched melanoma cells (E:T = 4:1) on monocultures of the same melanoma cells (**Supplementary Fig. S3A**), indicating that while TIL-mediated cancer cell killing requires physical contact between TIL and their target cells, soluble factors released during TIL-mediated cancer cell lysis are insufficient to induce cell death.

Since the scRNA-seq of expanded TIL (see Fig. 1D-F) confirmed the low abundance or absence of NKT, MAIT, and γδ T cells, we next examined if the expanded TIL expresses NK receptors (i.e., *NCR*, *NKG2*, and *KIR* receptors) and possess innate-like cytotoxic properties (33,34). However, expression of activating receptors and adhesion molecules, including *NCR1* (NKp46)*, NCR2* (NKp44), *NCR3* (NKp30), *KLRC2* (NKG2C), *KLRK1* (NKG2D), *CD56* (NCAM-1) (**Supplementary Fig. S3B-C**), and *KIR2DL1*, *KIR2DL3*, *KIR2DL4*, and *KIR3DL1* were low or absent in expanded TIL at either transcriptional or protein level (**Supplementary Fig. S3B**). Contact-dependent, pMHC-TCR independent killing of cancer cells by FAS-FASL and NKG2D-NKG2DL has previously been described (33,35). We then examined the possible role of FAS ligand (FASL, CD95L) interaction with FAS (CD95) on cancer cells (35). Single-cell RNA-seq analysis of 10049 melanoma cells (*n* = 5,519 cells) and surface staining confirmed that FAS is moderately expressed on melanoma cells (**Supplementary Fig. S3D-E)**, although FAS-neutralizing antibodies had no effect on cancer cell lysis by autologous TIL (**Fig. 4G, Supplementary Fig. S3F**). To further confirm this, we knocked out FAS in 4 different melanoma cell lines and performed autologous co-culture experiments at different ratios and over the time course of 3 days, which revealed that melanoma cells lacking FAS remain sensitive to TIL-mediated killing (**Supplementary Fig. S3G-J)**. Moreover, low level expression of NKG2DL (e.g., MICA/B) was observed in a minority of cancer cells (**Supplementary Fig. S3K-L**), and treatment with neutralizing antibodies against MICA/B, NKG2D, or both had no effect on TIL-mediated cancer cell lysis (**Fig. 2H**). In addition, recent studies showed that granzyme B- and perforin-mediated cytolytic function also require proximity between effector and target cells (36). However, small molecule inhibitors of granzyme B (Granzyme B inhibitor II) and perforin (Perforin-IN-2 (37)) had no effect on TIL-mediated cancer cell killing (**Fig. 2I, Supplementary Fig. S3M**). Taken together, these data demonstrated that TIL-mediated killing is contact-dependent but does not rely on unconventional T cells (e.g., NKT, MAIT, γδ T cells), and does not require FAS-FASL or NKG2D-NKG2DL interactions, or granzyme B or perforin-mediated cytolysis (**Fig. 2J**).

### TIL-mediated cancer cell killing requires LTBR and IFN sensing pathways

To nominate cancer cell-intrinsic genes and pathways critical for elimination by TIL, we performed a genome-scale, *in vitro* loss-of-function CRISPR screen using Cas9-expressing patient-derived melanoma cells (10049) co-cultured with autologous TIL (**Fig. 3A**). The quality control analyzes indicated excellent screen performance with the vast majority of single guide RNAs (sgRNAs) well-represented in all conditions (**Supplementary Fig. S4A-E**). To identify key regulators of TIL-mediated cancer cell killing, we compared sgRNA abundances between cells under selective pressure following TIL challenge (E:T = 3:1) compared to Output (E:T = 0:1) and confirmed that these sgRNAs are not scored in comparison between Output and Input (cell pools used for screen) (**Supplementary Fig. S4F**), such that negative scores signify enhanced sensitivity to TIL-mediated killing and positive scores signify resistance. The top scoring enriched sgRNAs included the lymphotoxin β receptor (*LTBR*), genes related to death receptor signaling (*FADD*, *CASP8*, *RIPK1*), and IFN sensing (*JAK1*, *JAK2*, *STAT1*, *IFNGR1*, *IFNGR2*) pathways (**Fig. 3B, Supplementary Fig. S4G-H**). Of note, other TNFR family members, including *TNFR1*, *FAS*, and *TNFSF10* (TRAIL), did not score as enriched sgRNAs in the screen (**Supplementary Fig. S4I**). Upon ligand binding, LTβR engagement can promote cancer cell death (10,13,38–40), including RIPK1-dependent cell death (41). The IFN-JAK/STAT signaling pathway has been well-known to induce cell cycle arrest and trigger cell death (42,43). We previously found the coordinated TNFR signaling and IFN sensing in immune-mediated cancer cell death (44). Supportingly, the top-scoring depleted sgRNAs included several gene targets involved in nuclear factor kappa-light-chain-enhancer of activated B cells (NF-κB) (*CHUK*, *OTULIN*, *CFLAR*, *RNF31*, *RELA*, *RELB*, *IKBKG*, *IKBKB*) and interferon (*PTPN2*) signaling pathways (**Fig. 3B, Supplementary Fig. S4G-H**) (42,45,46), underscoring the importance of these two pathways.

**Figure 3.**
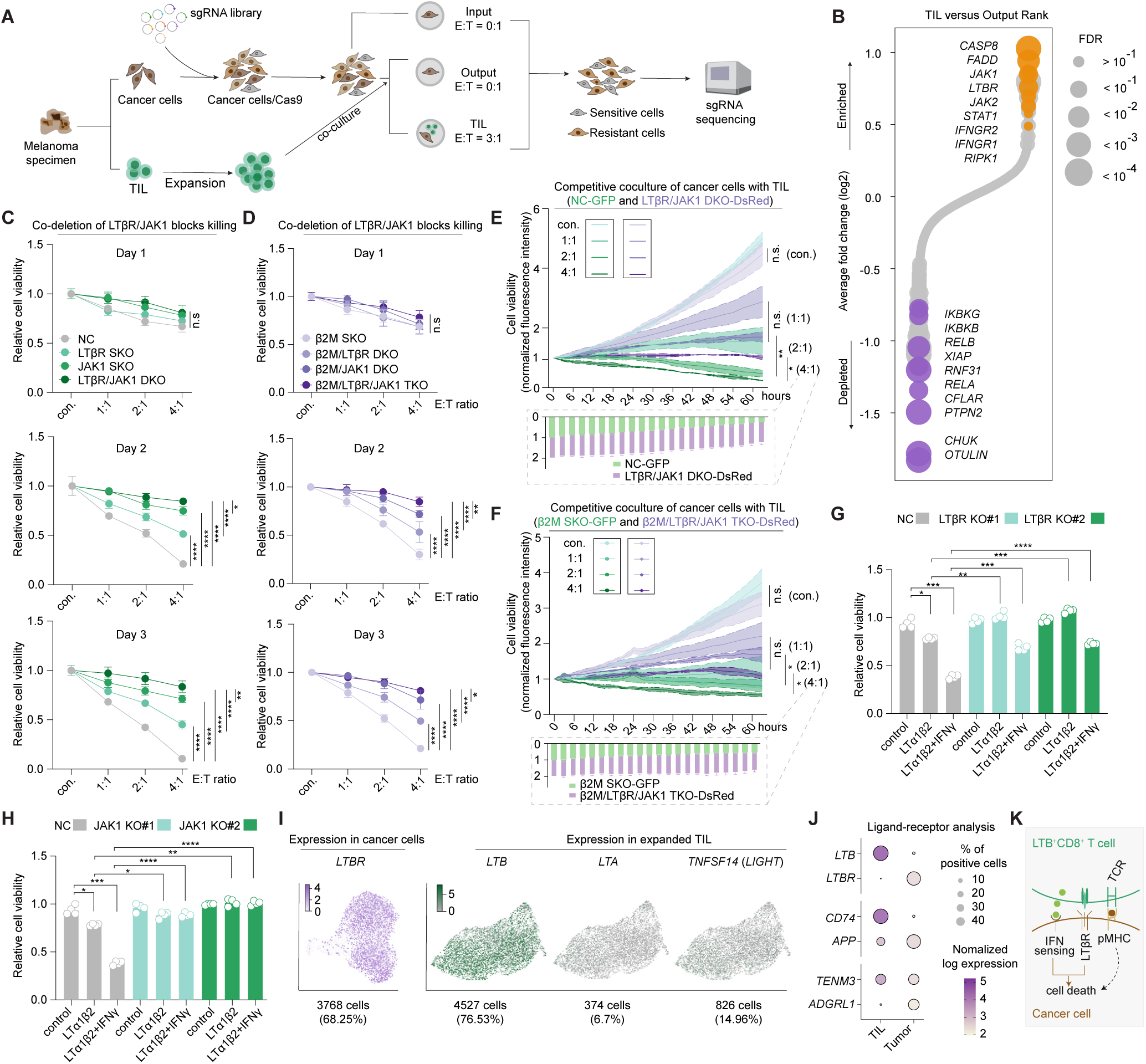
| TIL-mediated cancer cell killing requires LTBR and IFN sensing pathways. **A**, Schematic of genome-scale, loss-of-function CRISPR screen in cancer cells cocultured with autologous TIL. **B**, Genes ranked by TIL (E:T = 3:1) versus Output (E:T = 0:1)-normalized fold change for genome-scale screens, with circle size corresponding to FDR by STARS. Genes with 4-5 mapped guides are visualized; fold changes are calculated only from 4 top-performing guides per gene. Selected top genes with FDR values < = 0.3 at each end are listed and arranged by statistical significance. **C-D**, Viability assessment of the indicated melanoma (10049) cells after co-culture with autologous TIL compared with unstimulated cells. Mean values (circles) +/− s.d. (bars) are shown (*n* = 3; two-way ANOVA with Tukey’s multiple-comparison test). **E-F**, Competition assays using equal proportions of (**E**) control (GFP) and LTβR/JAK1 DKO (DsRed) or (**F**) *B2M*-null (GFP) and *B2M*-null/LTβR/JAK1 TKO (DsRed) 10049 melanoma cells co-cultured with autologous TIL at different E:T ratios at indicated times. Normalized fluorescence intensity of viable cancer cells (mean +/− s.d.; shaded region) is shown (*n* = 3, two-way ANOVA with Tukey correction of multiple comparisons). Inset stacked bar blot (E:T = 4:1) depicts proportional decrease in GFP and DsRed cells over time (*n* = 3, two-way ANOVA with Tukey correction of multiple comparisons). **G-H**, Viability assessment of (**G**) LTβR KO and (**H**) JAK1 KO melanoma cell lines (10049) treated with recombinant LTɑ1β2 (50 ng/mL) and/or IFNγ (40 ng/mL) for 2 days compared to untreated control. Individual values (open circles) are shown (*n* = 4; 2-sided paired *t-*test). **I**, UMAP embedding depicting the single-cell expression of indicated genes in 10049 TIL and cancer cells. **J**, Top 3 ligand-receptor pairs expressed in 10049 TIL and cancer cells determined by CellphoneDB. **K**, Schematic showing coactivation of LTβR and IFN sensing pathways can induce cancer cell death. \**P* < 0.05; \*\**P* < 0.01; \*\*\**P* < 0.001; **** *P* < 0.0001; *ns*, not significant.

### LTβR/IFN sensing is necessary and sufficient to induce cancer cell death

Next, we set out to confirm the role of LTβR and IFN sensing in TIL-mediated killing in cancer cells with or without class I HLA expression. Cancer cell-specific deletion of *LTBR* or *JAK1* alone provided modest protection against TIL-mediated killing, whereas co-deletion of *LTBR* and *JAK1* provided near complete rescue in both control and *B2M*-null melanoma cells (**Fig. 3C-D, Supplementary Fig. S4J-M**). These observations were further supported by time-lapse, image-based competitive killing assay in which DsRed-labeled melanoma cells lacking both *LTBR* and *JAK1* (sg*LTBR* and sg*JAK1*; LTβR/JAK1 DKO-DsRed) were protected from TIL-mediated killing whereas control melanoma cells (NC-GFP) were eliminated in a time- and dose-dependent manner (**Fig. 3E, Supplementary Fig. S5A-C**). DsRed-labeled melanoma cells lacking *B2M, LTBR*, and *JAK1* (sg*B2M*, sg*LTBR,* sg*JAK1*; β2M/LTβR/JAK1 TKO-DsRed) were similarly protected from TIL-mediated killing (**Fig. 3F, Supplementary Fig. S5A-B, 5D**). Consistent with these findings, pharmacologic inhibition of JAK1/2 (ruxolitinib) protected both control and *B2M*-null melanoma cells from TIL-mediated killing (**Supplementary Fig. S5E**).

Cancer cell-specific deletion of *CASP8* or *FADD* also protected melanoma cells from TIL-mediated killing (**Supplementary Fig. S5F-H**), as did pre-treatment with a pan-caspase (Q-VD-Oph) or caspase 8 (zIETD-fmk) selective inhibitor (**Supplementary Fig. S5I**). Pre-treatment with a RIPK1 inhibitor (Nec-1s) offered partial protection from TIL-mediated killing, whereas the pyroptosis inhibitor dimethyl fumarate (DMF) and the ferroptosis inhibitor ferrostatin-1 had no effect on TIL-mediated cancer cell lysis (**Supplementary Fig. S5I**). Combining caspase inhibition with necroptosis inhibition offered further protection against TIL-mediated killing **(Supplementary Fig. S5J**), suggesting that TIL-mediated killing requires both cell extrinsic apoptosis and necroptosis cell death pathways.

To determine if activation of LTβR and IFN signaling was sufficient to induce cancer cell death, we next examined cell viability of melanoma cells treated with recombinant LTβR ligands (LTɑ_1_β_2_ and LIGHT) with or without IFNγ. While LTβR activation with LTɑ_1_β_2_ or LIGHT resulted in a modest decrease in cell viability, a combination of either LTβR ligand with IFNγ dramatically enhanced cancer cell death that was abrogated by deletion of *LTBR* (**Fig. 3G, Supplementary Fig. S5K**), *JAK1* (**Fig. 3H, Supplementary Fig. S5L**), *JAK2* **(Supplementary Fig. S5M-O**), or *FADD* (**Supplementary Fig. S5P-Q**) in cancer cells. This suggests that recombinant LTβR ligands (LTɑ_1_β_2_ and LIGHT) and IFNγ are sufficient to induce cancer cells death and further demonstrates the role of FADD in LTβR- and IFN-driven cytotoxicity against cancer cells. Moreover, we confirmed patient-derived melanoma cell lines (*n* = 8) respond to recombinant LTβR ligands (LIGHT, LTɑ_1_β_2_) +/− IFNγ, observing a range of sensitivities (**Supplementary Fig. S6A-P**) despite similar levels of LTβR (**Supplementary Fig. S6Q**), suggesting that variable activation of tumor-intrinsic immune evasion pathways may limit cell death. Together, these findings demonstrated that coactivation of LTβR and IFN sensing pathways is both necessary and sufficient to induce cancer cell death.

### LTβR ligands expressed by expanded TIL

We next examined the expression of *LTBR* and LTβR ligands (*LTB*, *LTA*, *TNFSF14*) in melanoma cells and expanded TIL, respectively. Consistent with our flow cytometry data (see Supplementary Fig. S6Q), we observed high expression of *LTBR* in melanoma cells (**Fig. 3I**). High expression of *LTB* was observed in expanded TIL, whereas *LTA* and *TNFSF14* (encoding LIGHT) were expressed at low levels in a minority of TIL (**Fig. 3I**). Importantly, a subset of tumoricidal TIL was found to co-express *LTA* and *LTB* (*LTA*^+^*LTB*^+^) (**Supplementary Fig. S6R**). *IFNG* was also commonly expressed in *LTA*^+^*LTB*^+^ cells (*LTA*^+^*LTB*^+^*IFNG*^+^) (**Supplementary Fig. S6S**), and in TIL expressing LIGHT (*TNFSF14*^+^*IFNG*^+^) **(Supplementary Fig. S6T**), indicating that while *LTB* is broadly expressed, sub-populations of tumoricidal CD8^+^ TIL express all relevant effector molecules (*LTB*, *LTA*, *IFNG*, *TNFSF14*) necessary to activate LTβR and IFN sensing pathways in cancer cells. Further, ligand-receptor analysis using CellPhoneDB (47) identified LTB-LTBR as the top-scoring ligand-receptor pair (**Fig. 3J**). These findings further support a role for LTBR-LTB interactions in TIL-mediated cancer cell killing (**Fig. 3K**).

### *LTB^+^*CD8^+^ T cells are enriched in expanded TIL

Given the potent tumoricidal activity of expanded CD8^+^ TIL, we wondered if a relative paucity of *LTB*^+^CD8^+^ T cells or spatial restriction (e.g., immune exclusion) might preclude interaction with cancer cells in the native tumor microenvironment (TME). Thus, we performed multiplexed immunofluorescence with RNA *in situ* hybridization (RNA-ISH) of two melanoma specimens (10049 and 10222) to examine the extent of infiltration of *LTB*-expressing *CD8a*^+^ T cells relative to *LTBR*-expressing melanoma cells and their spatial organization in the TME. We observed a low level of *CD8a*^+^ infiltration (0.86% and 14.86%) and a very low proportion of *LTB*-expressing *CD8a*^+^ T cells relative to *LTBR*-expressing SOX10/S100^+^ melanoma cells (0.06% and 5.9%) (**Fig. 4A-B, Supplementary Fig. S7A-D**). Moreover, the average distance between *LTBR*^+^SOX10/S100^+^ melanoma cells and *LTB*^+^*CD8a*^+^ T cells was greater than 50 μm in the majority of interactions examined in the two specimens analyzed (**Fig. 4C, Supplementary Fig. S7E**). These findings confirmed that *LTB*^+^CD8^+^ T cells are present at low levels in the native TME, and that they are physically separated from *LTBR*^+^ melanoma cells.

**Figure 4.**
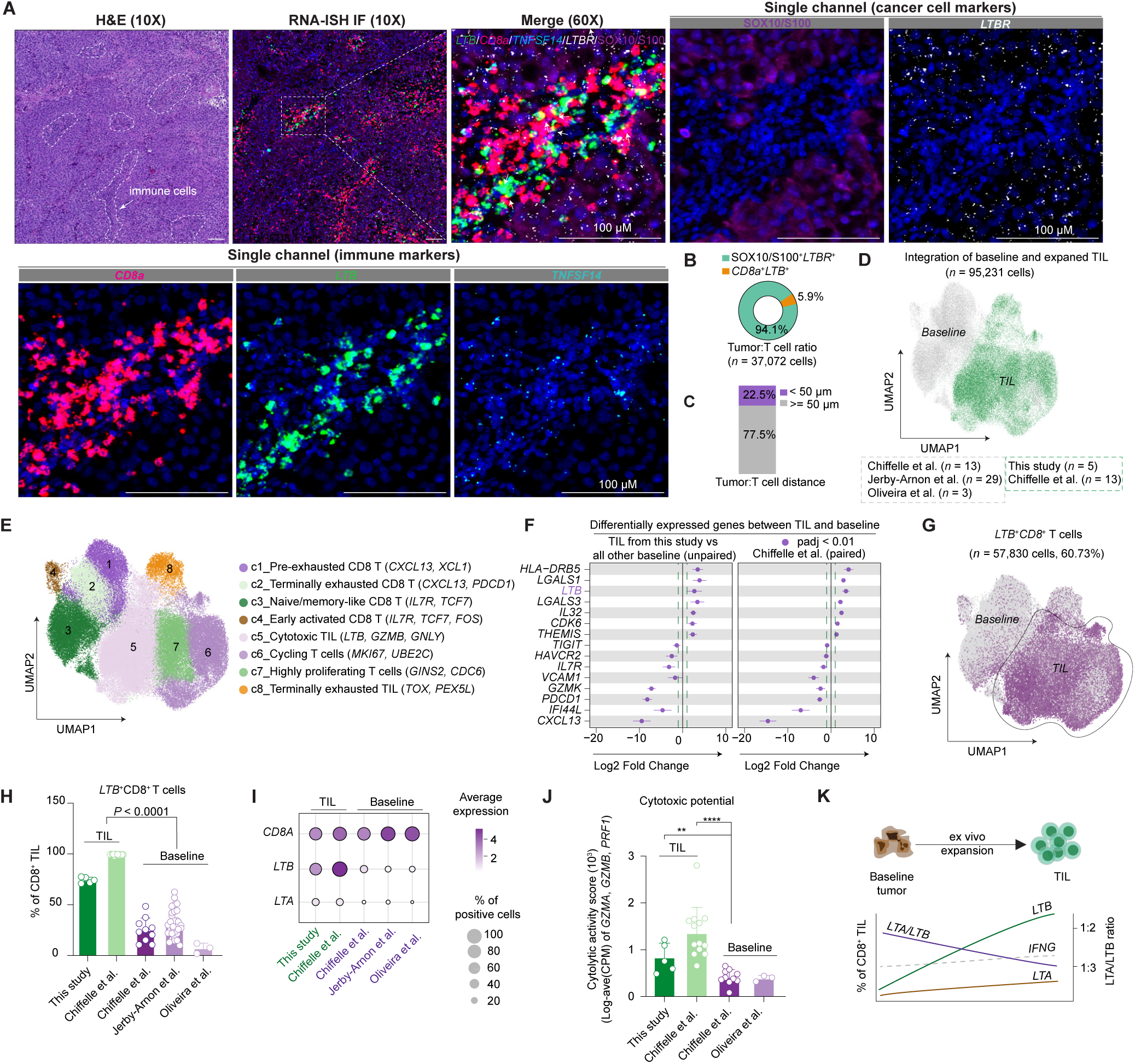
| *LTB^+^*CD8^+^ T cells are enriched in expanded TIL. **A**, Representative H&E (*upper left*) and immunofluorescence (IF) with RNA *in situ* hybridization (RNA-ISH) images (10X and 60X magnification) from excisional melanoma biopsies (10222) showing the expression of indicated genes and proteins. **B,** Percentage of SOX10/S100*^+^LTBR^+^* melanoma cells and *LTB*^+^*CD8a*^+^ T cells in cells analyzed. **C**, Percentage of SOX10/S100^+^*LTBR^+^* melanoma cells within the indicated distance of *LTB*^+^*CD8*^+^ T cells. **D-E**, UMAP projection of CD8^+^ T cells (*n* = 95,231 cells) from scRNA-seq analysis of (**D**) baseline tumor digests and expanded TIL from this study and published datasets for which (**E**) 8 unique CD8^+^ T cell clusters were identified. **F,** Selected differentially expressed genes between baseline tumor digests and expanded *CD8*^+^ TIL as indicated. **G,** UMAP embedding depicting the single-cell expression of *LTB* in integrated baseline tumor digests and expanded TIL. **H,** Percentage of *CD8*^+^ T cells belonging to *LTB* high clusters in baseline tumor digests and expanded TIL across different datasets (individual open circle represents one sample, 2-sided unpaired *t*-test). **I,** Dot plot showing the expression of selected genes among baseline and expanded TIL. **J**, Cytolytic activity score of TIL calculated by the log-average (CPM) of *GZMA, GZMB,* and *PRF1* in baseline tumor digests and expanded TIL across different datasets (individual open circle represents one sample, 2-sided unpaired *t*-test). \*\**P* < 0.01; **** *P* < 0.0001. **K,** Schematic showing the dynamic change of *LTA*, *LTB,* and *IFNG* expression as well as *LTA*/*LTB* ratio during TIL expansion.

Given the abundance of expanded *LTB*^+^ CD8^+^ TIL despite the paucity of *LTB*^+^ CD8^+^ T cells in baseline tumor specimens, we reasoned that *LTB*^+^ CD8^+^ T cells might be enriched during TIL expansion. To examine this further, we performed an integrated scRNA-seq analysis (*n* = 95,231 cells) using expanded TIL (*n* = 5) from this study together with published scRNA-seq datasets of expanded TIL (*n* = 13) and baseline tumor digests (*n* = 45) from melanoma patients (5,48–50). We aggregated datasets with a focus on *CD8*^+^ T cells and performed clustering in order to create a stable uniform manifold approximation and projection (UMAP) embedding (**Fig. 4D-E)**. Interestingly, *CD8*^+^ T cells from baseline tumor digests and expanded TIL occupied distinct clusters, with expanded TIL enriched in clusters 5 (cytotoxic), 6 (cycling), and 7 (highly proliferating), whereas clusters 1 (pre-exhausted), 2 (terminally exhausted), and 3 (naive/memory-like) were observed in baseline tumor digests (**Fig. 4D-E**, **Supplementary Fig. S7F-G**). Differential gene expression analysis confirmed upregulation of *LTB* in expanded TIL, whereas higher levels of *CXCL13* and *PDCD1* were observed in CD8^+^ T cells from baseline tumor digests (**Fig. 4F**). A higher proportion of *LTB*^+^*CD8*^+^ cells and increased expression of *LTB* and *LTA* were observed in expanded TIL relative to baseline tumor digests (**Fig. 4G-I**), as was a gene expression-based cytotoxicity score(29) (**Fig. 4J**). Taken together, these findings confirmed dramatic transcriptional differences between expanded TIL and *CD8*^+^ T cells from baseline tumor digests, and demonstrated that *LTB*^+^*CD8*^+^ T cells are enriched in expanded TIL (**Fig. 4K**).

### Dynamic regulation of lymphotoxin in TIL-melanoma cocultures

We next examined the expression of lymphotoxin (*LTA* and *LTB*) and *IFNG* in the expanded TIL before and after co-culture with cancer cells (E:T of 4:1, 24 hours). TIL were collected pre- and post-coculture with autologous control (4 pairs) or *B2M*-null (2 pairs) cancer cells for scRNA-seq analysis (**Fig. 5A, Supplementary Fig. S7H-I**). While negligible changes in cluster enrichment and TCR clonality were observed before and after coculture with cancer cells (**Supplementary Fig. S7J-L**), we observed dynamic changes in *LTA* expression and the ratio of LTA/LTB expression (**Fig. 5B-D, Supplementary Fig. S7M-P**). Expression of *LTA* was minimal at baseline, but was significantly increased in TIL following coculture with control and *B2M*-null melanoma cells (**Fig. 5B-D, Supplementary Fig. S7M-P**). Interestingly, high baseline expression of *LTB*, *GZMB*, and *IFNG* was observed in expanded TIL which was preserved in TIL following coculture with control and *B2M*-null melanoma cells (**Fig. 5C, Supplementary Fig. S7O**).

**Figure 5.**
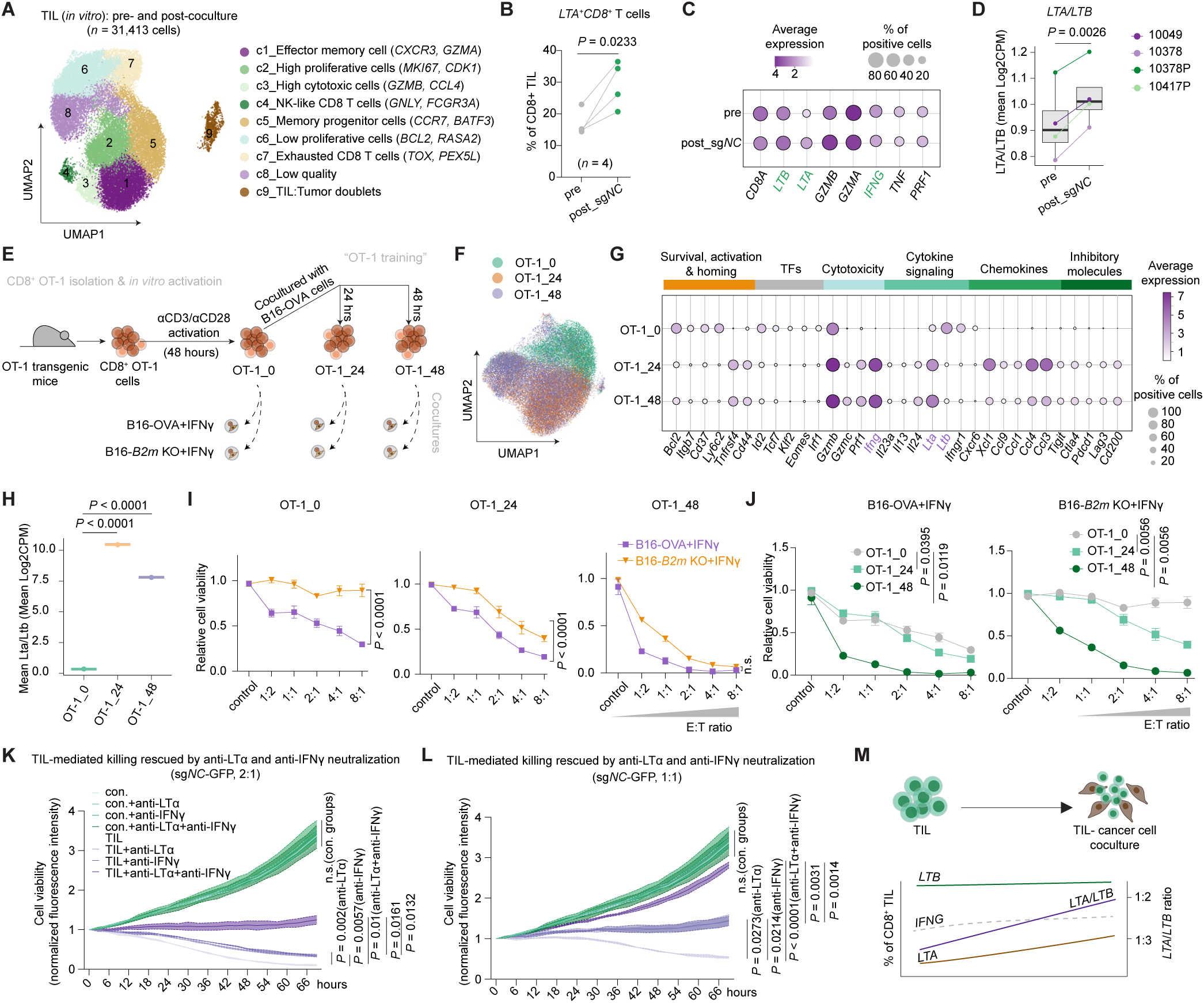
| Dynamic regulation of lymphotoxin in TIL-melanoma cocultures. **A**, UMAP projection of expanded CD8^+^ TIL pre- and post-coculture with cancer cells for 24 hours (*n* = 31,413 cells) with 9 unique populations identified from scRNA-seq analysis. **B**, Percentage of *LTA*^+^*CD8*^+^ TIL before and after coculture with control (sg*NC*) melanoma cells. Individual open circles represent one TIL (*n* = 4; 2-sided paired *t*-test). **C**, Dot plot showing expression of selected genes before and after coculture with control (sg*NC*) melanoma cells. **D**, Bar plot examining the log2 fold change (mean CPM) of the *LTA*/*LTB* ratio in TIL before and after coculture with sg*NC*. **E**, Schematic of experimental design. CD8^+^ OT-1 T cells were isolated from the spleens of OT-1 transgenic mice, activated by αCD3 and αCD28 antibodies, and expanded *in vitro* with IL-2 (50 ng/mL) for 48 hours. Activated OT-1 T cells were then co-cultured for 0, 24, and 48 hours with B16-OVA cells pretreated with IFNγ (E:T = 8:1) and named as OT-1_0, OT-1_24, and OT-1_48 after dead cell removal. Next, these OT-1 cells were cocultured with B16-OVA or B16-*B2m* KO cells pretreated with IFNγ, respectively. **F**, UMAP embedding of scRNA-seq data for OT-1_0, OT-1_24, and OT-1_48 cells. **G,** Dot plot showing the expression of selected genes among OT-1_0, OT-1_24, and OT-1_48 cells. **H**, Bar plot examining the log2 fold change (mean CPM) of *Lta*/*Ltb* ratio in OT-1_0, OT-1_24, and OT-1_48 cells. **I-J**, Cell viability of B16-OVA+IFNγ and B16 *B2m* KO+IFNγ cocultured with OT-1_0, OT-1_24, and OT-1_48 cells. Mean +/− s.d. (bar) is shown (*n* = 3; Two-way ANOVA with Tukey correction of multiple comparisons). **K-L,** Normalized fluorescence intensity of viable melanoma cells (10049, sg*NC*-GFP) cocultured with autologous TIL at the E:T ratio of (**K**) 2:1 and (**L**) 1:1 with or without the indicated neutralizing antibodies (anti-LTα, 10 μg/mL; anti-IFNγ, 10 μg/mL) over indicated time points. Mean +/− s.d. (shaded region) is shown (*n* = 3, two-way ANOVA with Tukey correction of multiple comparisons; *ns*, not significant). **M,** Schematic showing the dynamic change of *LTA*, *LTB,* and *IFNG* expression as well as *LTA*/*LTB* ratio upon coculture with cancer cells.

We next sought to determine if the dynamic change of lymphotoxin upon TIL-cancer cell coculture is conserved in an established antigen-specific T cell co-culture system. We first examined the differential gene expressions of TCR transgenic OT-1 CD8^+^ T cells (specific for the ovalbumin-derived SIINFEKL peptide) after *in vitro* activation with ɑCD3/ɑCD28 (OT-1_0) and those further cocultured with ovalbumin-express B16 murine melanoma cells (B16-OVA) pretreated with IFNγ (resulting in the increase of H-2Kb, **Supplementary Fig. S8A**) for 24 hours (OT-1_24) and 48 hours (OT-1_48) (**Fig. 5E**). Unbiased clustering of scRNA-seq data from OT-1_0, OT-1_24, and OT-1_48 (*n* = 56,196 cells) revealed two dominant clusters, one occupied dominantly by OT-1_0 and the second by OT-1_24 and OT-1_48 cells (**Fig. 5F-G**). Analysis of differentially expressed genes revealed that OT-1_0 cells were more “stem-like”, with a higher expression level of *Tcf7* and *Ltb*, whereas OT-1_24 and OT-1_48 cells exhibited higher expression of effector cytokines and chemokines (*Lta*, *Ifng*), cytolytic molecules (*Prf1*, *Gzmb*, *Gzmc*), as well as co-inhibitory receptors (*Ctla4*, *Pdcd1*, *Lag3*, *Tigit*) (**Fig. 5G, Supplementary Fig. S8B**). These findings confirmed that dynamic regulation of *Lta* and *Ltb* is conserved in OT-1 after coculture with B16-OVA melanoma cells (**Fig. 5H**).

Next, we sought to determine if OT-1_24 or OT-1_48 acquired the capacity to eliminate B16-OVA cancer cells independent of class I MHC. As expected, OT-1_0 CD8^+^ T cells were unable to eradicate *B2m*-null B16 melanoma cells (lacking class I MHC surface expression) despite effectively eliminating control B16-OVA melanoma cells **(Fig. 5I-J)**. In contrast, OT-1_24 and OT-1_48 cells were able to kill both control B16-OVA and *B2m*-null B16 melanoma cells in a dose-dependent manner particularly at higher E:T ratios. Notably, OT-1_48 exhibited more potent activity against both control B16-OVA and *B2m*-null B16 melanoma cells compared to OT-1_24 (**Fig. 5I-J**), indicating that a longer duration of OT-1 co-culture with control B16-OVA (with intact class I MHC expression) results in enhanced tumoricidal capacity against both class I MHC-proficient (control) and -deficient (*B2m*-null) B16 melanoma cells. Taken together, these results confirmed dynamic regulation of lymphotoxin in an established antigen-specific T cell co-culture system, and revealed that antigen-specific CD8^+^ T cells can acquire the capability to kill cancer cells independent of MHC-I, though only after initial class I MHC-dependent killing.

Lastly, we examined the effect of neutralization of LTα +/− IFNγ on cancer cell killing mediated by human TIL. Anti-LTα (pateclizumab) in combination with anti-IFNγ rescued control and *B2M*-null melanoma cells from TIL-mediated killing to a greater extent than anti-LTα or anti-IFNγ alone (**Fig. 5K-L, Supplementary Fig. S8C-D**). Thus, while *LTB* expression is increased in expanded CD8^+^ TIL relative to baseline CD8^+^ TIL (see Fig. 4H-J), *LTA* expression is further upregulated following cancer cell encounter. This may represent a regulatory mechanism promoting formation of the LTα1β2 heterotrimer, which, together with other cytotoxic pathways (e.g., IFN), coordinates the function of tumoricidal *LTB*⁺ CD8⁺ TIL (**Fig. 5M**).

### *LTB*^+^*CD8*^+^ T cells are enriched in TIL products of patients responsive to lifileucel

While our findings established a role for LTβR and IFN sensing pathways in TIL-mediated cancer cell killing, as well as the dynamic regulation of lymphotoxin in TIL, the clinical significance of these observations remained uncertain. Thus, we set out to determine if enrichment of *LTB^+^CD8*^+^ T cells was associated with clinical response to TIL cell therapy in patients with advanced ICB-refractory melanoma (C-144-01 Study, ClinicalTrials.gov ID: NCT02360579) (51–53). First, we examined scRNA-seq data from TIL products (lifileucel; *n* = 34) for patients with advanced melanoma, generating a UMAP embedding (*n* = 116,540 cells) with 15 sub-clusters of *CD8*^+^ TIL (**Fig. 6A-B, Supplementary Fig. S9A**). Next, we examined the expression of *LTB* in the lifileucel embedding and observed marked enrichment of *LTB*^+^*CD8*^+^ TIL in cluster 0 (hereafter referred to as “TIL-LTB^hi^_c0”) (**Fig. 6C**). Enrichment of TIL-LTB^hi^_c0 was observed in patients experiencing a clinical response (R) or stable disease (SD) to lifileucel treatment compared to those with progressive disease (PD) (**Fig. 6D**). Of note, the proportion of *LTB*^+^ TIL from all CD8^+^ clusters was significantly lower in patients with PD compared to patients with R or SD to lifileucel **(Fig. 6E**). Gene set scoring analysis of TIL-LTB^hi^_c0 compared to other lifileucel clusters (1–14) revealed a higher score for stem-cell like and cytotoxicity signatures (**Fig. 6F-G**) and a lower score for a signature associated with terminal differentiation (6) (**Fig. 6H**). These findings demonstrate that the abundance of tumoricidal *LTB*^+^*CD8*^+^ in expanded TIL is associated with clinical benefit to TIL cell therapy with lifileucel.

**Figure 6.**
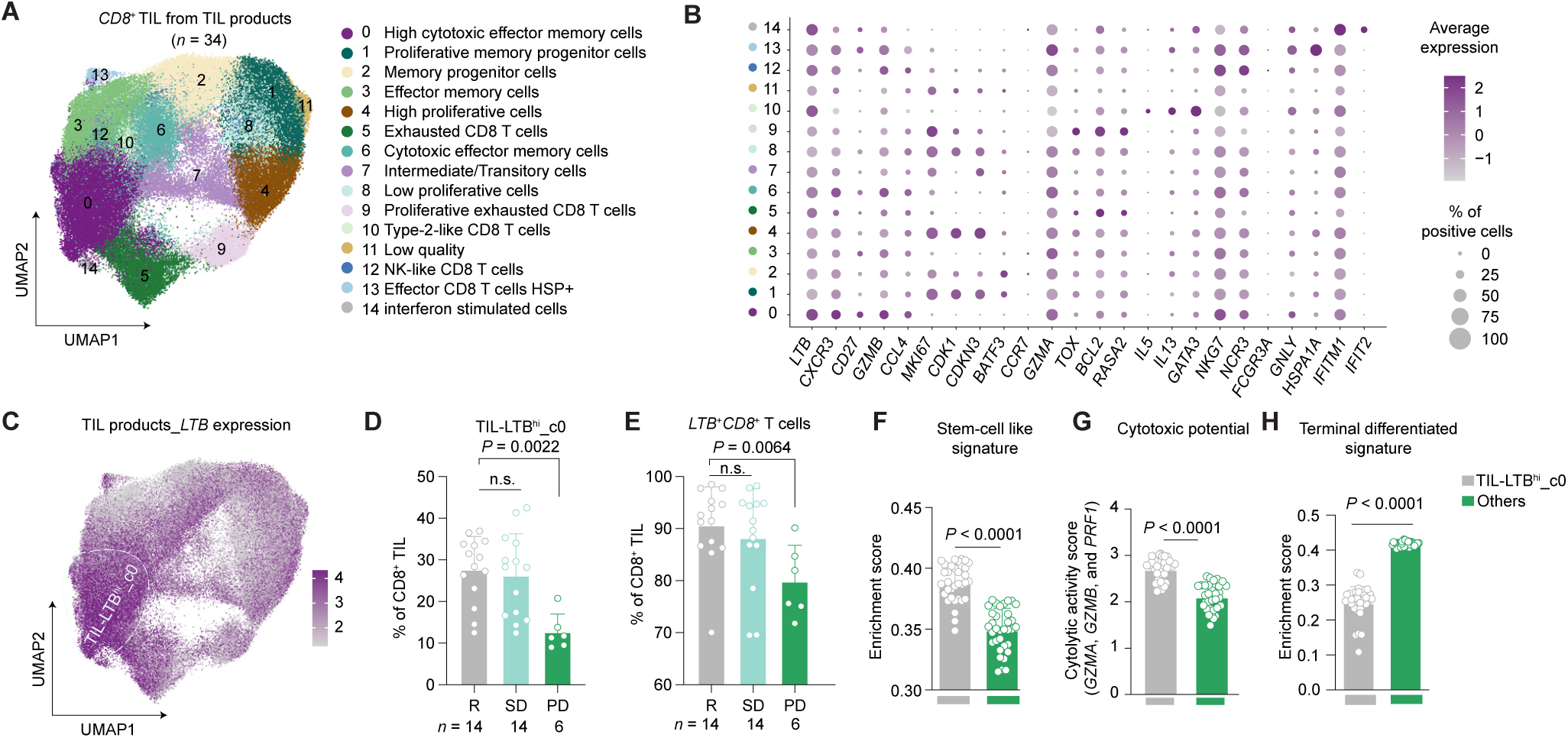
| *LTB*^+^*CD8*^+^ T cells are enriched in TIL products of patients responsive to lifileucel. **A**, UMAP embedding of scRNA-seq data from CD8^+^ TIL (*n* = 116,540 cells) from lifileucel TIL products (*n* = 34) identifying 15 unique clusters. **B,** Dot plot showing the representative marker genes that are differentially expressed across different clusters. **C**, UMAP projection of *LTB* expression in lifileucel TIL products. **D-E**, Bar plots showing the percentage of (**D**) TIL-LTB^hi^_c0 and (**E**) *LTB*^+^*CD8*^+^ TIL of all CD8^+^ TIL grouped by best overall response (BOR); responder (R), stable disease (SD), or progressive disease (PD). Patient-level pseudobulk profiles were generated from the scRNA-seq data and gene set scoring was performed as described in the methods. Mean +/− s.d. (bars) and individual values (open circles) are shown (Kruskal-Wallis test with Dunn’s multiple comparisons test; *ns*, not significant). **F-H**, Bar plots showing (**F**) stem-like signature enrichment scores, (**G**) cytolytic activity scores, and (**H**) terminal differentiation enrichment scores for LTB^hi^ cluster 0 (TIL-LTB^hi^_c0) compared to all other CD8^+^ TIL clusters. Mean +/− s.d. (bars) and individual values (open circles) are shown (Kruskal-Wallis test with Dunn’s multiple comparisons test).

### *LTB*^+^CD8^+^ TIL are expanded from putative neo-antigen reactive CD8^+^ T cells

We next sought to determine if tumoricidal *LTB*^+^*CD8*^+^ TIL arise from *LTB*^+^*CD8*^+^ T cells in tumor digests from resection specimens used for TIL expansion. To address this question, we performed matched scRNA-seq and scTCR-seq analysis from baseline tumor digests and TIL products (*n* = 7). We then did unbiased clustering on *CD8*^+^ T cells (*n* = 4,401) in baseline tumor digests identifying 10 distinct clusters (**Fig. 7A, Supplementary Fig. S9B**). Next, we projected *LTB* expression onto the embedding generated from tumor digests and observed enrichment of *LTB*^+^*CD8*^+^ TIL (hereafter referred to as “baseline-LTB^hi^”) in cluster 0 (recently activated), 4 (central memory), 5 (naive), and 7 (short-lived effector) (**Fig. 7B**). Evaluation of differentially expressed genes revealed upregulation of *LTB* in expanded TIL, and preferential expression of *CXCL13* and *PDCD1* in baseline tumor digests (**Fig. 7C**). Consistent with these observations, *LTB*^+^*CD8*^+^ T cells were enriched in expanded TIL (lifileucel) compared to baseline tumor digests (**Fig. 7D**).

**Figure 7.**
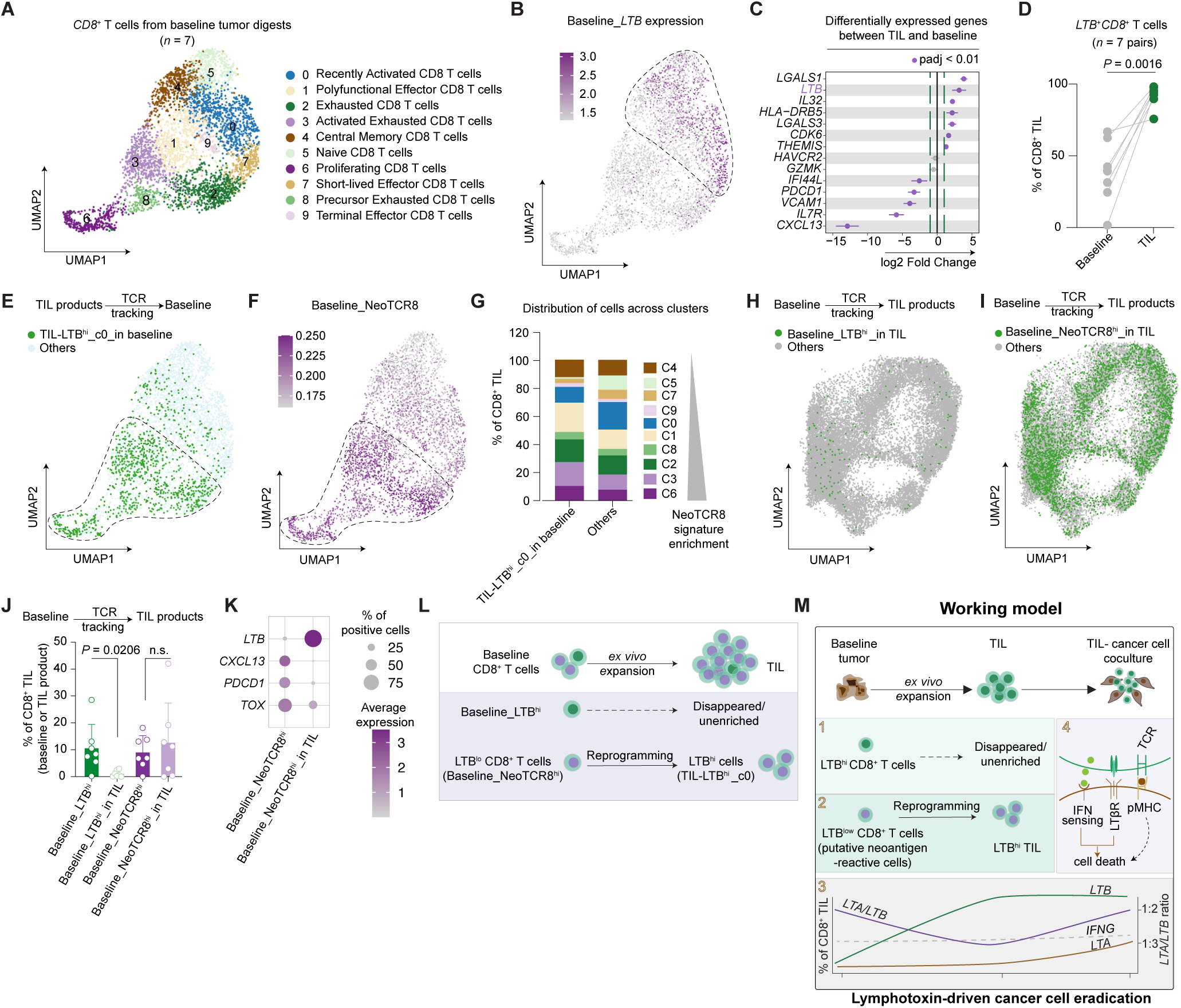
| *LTB*^+^CD8^+^ TIL are expanded from putative neo-antigen reactive CD8^+^ T cells. **A**, UMAP embedding of scRNA-seq data from CD8^+^ T cells (*n* = 4,401 cells) in baseline tumor digests (*n* = 7). **B**, UMAP projection of *LTB* expression in baseline tumor digests. **C**, Selected differentially expressed genes between baseline tumor digests and TIL product as indicated. **D,** Percentage of *LTB*^+^*CD8*^+^ cells in TIL products compared to matched baseline tumor digests (*n* = 7, 2-sided paired *t*-test). **E**, Projection of TCRs shared with TIL-LTB^hi^_c0 (green) compared to TCRs from all other CD8^+^ TIL clusters (light blue) on the UMAP embedding of CD8^+^ T cells of the baseline tumor digests. **F**, NeoTCR8 signature expression projected on the UMAP embedding of CD8^+^ T cells of the baseline tumor digests. **G**, Proportions of CD8^+^ T cell clusters in baseline tumor digests with TCRs shared with TIL-LTB^hi^_c0 compared to all other CD8^+^ TIL clusters (ordered by NeoTCR8 signature enrichment). **H-I**, Projection of TCRs shared with (**H**) Baseline_LTB^hi^ and (**I**) Baseline_NeoTCR8^hi^ CD8^+^ T cells on the UMAP embedding from CD8^+^ TIL (lifileucel TIL products). **J**, Bar plot showing the percentage of cells expressing the LTB^hi^ or NeoTCR8^hi^ signature in baseline tumor digests, and the percentage of cells expressing the same TCRs in lifileucel products. Mean +/− s.d. (bars) and individual values (open circles) are shown (*n* = 7, 2-sided paired *t*-test, *ns*, not significant). **K**, Dot plot showing expression of indicated genes in NeoTCR8^hi^ CD8^+^ T cells in baseline tumor digests compared to CD8^+^ TIL in lifileucel with shared TCRs. **L**, Schematic showing the dynamic change of LTB^hi^ and NeoTCR8^hi^ (also LTB^lo^) CD8^+^ T cells during expansion. **M**, Schematic showing the proposed working model.

To identify the clonal origins of TIL-LTB^hi^_c0, we leveraged paired scTCRseq data to track the TCR clones of expanded CD8^+^ TIL (TIL product) back to baseline CD8^+^ TIL from tumor digests (TIL-LTB^hi^_c0_in baseline) (**Fig. 7E**). TCR backtracking confirmed that TIL-LTB^hi^_c0 arose from clusters 1 (polyfunctional), 2 (exhausted), 3 (activated exhausted), 6 (proliferating), and 8 (precursor exhausted) CD8^+^ TIL from tumor digests (**Fig. 7E and 7A**). Interestingly, these two populations (baseline-LTB^hi^ and TIL-LTB^hi^_c0_in baseline) appeared mutually exclusive, indicating that *LTB*^+^*CD8*^+^ TIL enriched in TIL-LTB^hi^_c0 did not arise from baseline-LTB^hi^ TIL (**Fig. 7E and 7B**). Lowery and colleagues previously described NeoTCR8, a 243-gene signature expressed by putative neoantigen-reactive CD8^+^ TIL that predicted antitumor, neoantigen-specific TCRs based on transcriptomic profiles (54). To determine if TIL-LTB^hi^_c0 cells were derived from putative neoantigen-reactive CD8^+^ T cells, we projected the NeoTCR8 signature on the baseline embedding (hereafter referred to as “baseline_NeoTCR8^hi^”) (**Fig. 7F**). We observed significant overlap between baseline_NeoTCR8^hi^ T cells and TIL-LTB^hi^_c0_in baseline T cells, indicating *LTB*^+^*CD8*^+^ TIL (TIL-LTB^hi^_c0) shared TCRs with putative neoantigen-reactive *CD8*^+^ T cells found in baseline tumor digests (baseline_NeoTCR8^hi^) (**Fig. 7E-G**). These data demonstrate that expanded CD8^+^ TIL enriched in TIL-LTB^hi^_c0 arise from putative neoantigen-reactive *CD8*^+^ T cells with low or absent *LTB* expression (baseline_NeoTCR8^hi^ or baseline-LTB^lo^ cells).

To examine the fate of cells enriched in baseline-LTB^hi^, we traced TCRs of these cells and projected them onto the embedding generated from TIL products (see Fig. 4a). Projection of TCRs from baseline_LTB^hi^ (Baseline_LTB^hi^) confirmed poor representation of T cell clonotypes in the expanded TIL product (Baseline_LTB^hi^_in TIL) (**Fig. 7H**). In contrast, CD8^+^ T cells in baseline tumor digests with high NeoTCR8 expression (Baseline_NeoTCR8^hi^) remain enriched in the TIL products (Baseline_NeoTCR8^hi^_in TIL) (**Fig. 7H-J**). Moreover, differential gene analysis between putative neoantigen-reactive *CD8*^+^ T cells in baseline tumor digests (Baseline_NeoTCR8^hi^) and their counterparts in TIL products (Baseline_NeoTCR8^hi^_in TIL) suggested Baseline_NeoTCR8^hi^_in TIL underwent dramatic transcriptional changes, with marked upregulation of *LTB* and concomitant downregulation of *CXCL13*, *PDCD1*, and *TOX* (**Fig. 7K**). Taken together, these results indicate that tumoricidal CD8^+^ TIL are not related to baseline-LTB^hi^ CD8^+^ T cells found in tumors, and that *LTB*^hi^ tumoricidal CD8^+^ T cells in TIL products arise from putative neoantigen-reactive *CD8*^+^ T cells with low or absent *LTB* expression (baseline-NeoTCR8^hi^; baseline_LTB^lo^) (**Fig. 7L-M**).

## DISCUSSION

Using patient-derived TIL-melanoma co-cultures we have identified and characterized a novel, therapeutically relevant subset of CD8^+^ TIL capable of eliminating cancer cells *in vitro* with or without canonical pMHC-TCR interactions. Elimination of cancer cells by these tumoricidal CD8^+^ TIL requires dual activation of LTβR and IFN sensing pathways, which is necessary and sufficient to induce cancer cell lysis. Tumoricidal CD8^+^ TIL express high levels of *LTB* and upregulate *LTA* upon coculture with cancer cells. Higher proportions of *LTB*^+^*CD8*^+^ T cells were observed in the TIL products of patients who experienced a clinical response to TIL therapy, and *LTB*^+^*CD8*^+^ TIL (TIL-LTB^hi^_c0) arise from putative neoantigen-reactive *CD8*^+^ T cells (with low *LTB* expression) in baseline tumor digests (baseline_NeoTCR8^hi^) (**Fig. 7M**).

Lifileucel, the first FDA-approved TIL product, has demonstrated a response rate of 31.4% in patients with advanced melanoma that has progressed after front-line treatment with ICB (and MAPK targeted therapy, for eligible patients) (53,55,56). The clinical success of TIL therapy for patients with ICB-refractory melanoma implies the existence of intratumoral, tumor-specific CD8^+^ T cells that were insufficiently stimulated in response to ICB treatment, but are expanded *in vitro* giving rise to tumoricidal CD8^+^ TIL. Here, we demonstrate that *LTB*^+^*CD8*^+^ T cells are enriched in expanded TIL, and higher proportions of *LTB*^+^*CD8*^+^ T cells (i.e., TIL-LTB^hi^_c0) in TIL products were observed in melanoma patients that achieved clinical response or stable disease following treatment with lifileucel. Whereas enrichment of stem-like CD8^+^ T cells has previously been associated with response to TIL therapy (6), our findings demonstrate that antitumor, putative neoantigen-reactive T cells in resected tumors undergo transcriptional reprogramming following *in vitro* expansion in the presence of IL-2 resulting in increased expression of *LTB* and cytotoxic gene signatures in addition to stem-like gene programs. These findings suggest that the ideal specimen for TIL harvest is one enriched for intratumoral neoantigen-reactive CD8^+^ T cell clonotypes with features of exhaustion that give rise to *LTB*-expressing CD8^+^ T cells. As such, strategies to improve the quality of expanded TIL products may require enhancements to (a) identification and isolation of neoantigen-reactive CD8^+^ T cells (e.g., high CXCL13), and (b) optimization of *in vitro* expansion protocols to increase the proportion of *LTB*-expressing CD8^+^ T cells may improve TIL products.

Quite unexpectedly, we also observed efficient *in vitro* TIL-mediated cancer cell elimination in the absence of pMHC-TCR interactions. We demonstrated that pMHC-TCR interactions were dispensable for *in vitro* TIL-mediated killing of matched melanoma cells using both anti-MHC-I neutralizing antibodies and CRISPR deletion of *B2M* and *TRAC*/*TRBC* in melanoma cells and TIL, respectively. The observation that *in vitro* expanded CD8^+^ TIL are capable of pMHC-TCR independent killing of cancer cells raises several critical questions about the precise role and requirement of pMHC-TCR in the recognition and elimination of malignant cells, as well as the timing of such interactions. Given the observed enrichment of putative neoantigen-reactive *CD8*^+^ T cell clonotypes in commercial TIL products, it is likely that pMHC-TCR interactions were critical for the formation of tumor-reactive (NeoTCR-hi) CD8^+^ T cells within the tumor (and/or tumor-draining lymph node). In the setting of chronic antigen exposure and an immune suppressive tumor microenvironment, these tumor-reactive CD8^+^ T cells likely became dysfunctional with diminished proliferative capacity, upregulation of co-inhibitory receptors, and reduced cytotoxic capacity. Following *in vitro* expansion in the presence of IL-2, tumor-reactive (NeoTCR-hi) CD8^+^ T cells underwent transcriptional reprogramming characterized by upregulation of *LTB* and signatures of stem-like properties and cytotoxic potential, with downregulation of signatures associated with T cell exhaustion. As a consequence, these tumor-reactive CD8^+^ T cells acquired the ability to eliminate tumor cells in a pMHC-TCR independent manner. While our observations with expanded, polyclonal TIL and matched melanoma cells cannot address the precise role and timing of pMHC-TCR interactions in the generation of tumor-reactive (NeoTCR-hi) CD8^+^ T cells, our observations with OT-1 T cells cocultures with B16-OVA melanoma cells suggests that pMHC-TCR interactions are requisite for the acquisition of tumoricidal properties. While recognition and elimination of B16-OVA by OT-1 T cells requires intact pMHC-TCR interactions initially, OT-1 dynamically acquire class I MHC independence following ‘training’ on control B16-OVA cells, during which upregulation of *Lta* and *Ifng* is observed in OT-I CD8⁺ T cells. Taken together, our observations support a model in which class I MHC-recognition of tumor-associated neoantigens gives rise to reversibly exhausted, tumor-reactive CD8^+^ T cells that are subsequently endowed with the capacity to eradicate cancer cells dependent on LTβR and IFN sensing, but independent of canonical pMHC-TCR interactions, following *in vitro* expansion and transcriptional reprogramming.

Coactivation of LTβR and IFN-JAK-STAT signaling pathways was necessary for TIL-mediated killing in both control and *B2M*-null cancer cells via a hybrid form of cell death reliant on both extrinsic apoptotic (e.g., *CASP8*, *FADD*) and necroptotic (e.g. *RIPK1*) signaling pathways. Additionally, we observed a range of sensitivities to both TIL- and cytokine-mediated cell death in patient-derived melanoma cell lines despite comparable expression of LTβR, suggesting that additional tumor-intrinsic factors may influence sensitivity to TIL therapy. Beyond these tumor-intrinsic factors, the dramatic upregulation of *LTB* in tumoricidal CD8^+^ TIL following *in vitro* expansion relative to native tumor-infiltrating CD8^+^ T cells, and the temporal transcriptional regulation of *LTA* and other effector molecules (e.g., IFNγ) following cancer cell engagement, indicates that TIL efficacy is also regulated dynamically by T cell-intrinsic factors. In addition to *LTB*, *LTA*, and *IFNG*, tumoricidal CD8^+^ TIL express high transcript levels of cytolytic molecules (e.g., *GMZB*, *PRF1*), though our results did not support a dominant role for granzyme B or perforin in TIL-mediated cancer cell killing.

These findings not only provide mechanistic insights into the role of LTβR/IFN signaling in class I MHC-independent, TIL-mediated cancer cell killing, but may also inform novel strategies to enrich specific TIL subsets during expansion or to augment specific TIL function(s). Further, insights from our study may facilitate the development of rational therapeutic strategies to leverage LTβR and IFN sensing pathways to recapitulate and enhance the tumoricidal properties of TIL for patients with advanced melanoma, and potentially other malignancies.

## METHODS

### Patient samples

Tumor samples were collected and analyzed according to Dana-Farber/Harvard Cancer Center IRB-approved protocols (DF/HCC 11-181). A cohort of patients (**Supplementary Table S1**) treated at Massachusetts General Hospital was processed, isolated, and *in vitro* expanded for generating patient-derived cell lines and TIL. These studies were conducted according to the Declaration of Helsinki and approved by the DF/HCC IRB. Response to treatment was determined radiographically, as previously described(44).

### C-144-01 study (ClinicalTrials.gov ID: NCT02360579)

The study was approved by the Institutional Review Board (IRB) at each site and was conducted in accordance with the Declaration of Helsinki and Good Clinical Practice guidelines of the International Conference on Harmonization. All patients provided written informed consent. The average age of the sub-data set of patients presented in this manuscript was 54.5 years old (s.d. +/− 14.0), with 35.3% female and 64.7% male.

### Animal treatment

The designs of animal studies and procedures were approved by the Broad Institute and Massachusetts General Hospital IACUC committees. Ethical compliance with IACUC protocols and institute standards was maintained. Specific pathogen-free facilities were used for the storage and care of all mice. CO_2_ inhalation was used to euthanize mice.

### CoMut plots

Graphs were generated based on Crowdy et al.(57), instructions can be found in Github https://github.com/vanallenlab/comut/blob/master/examples/documentation.ipynb. Python v3.9.13 was used to generate the plots.

### Cell lines

Patient-derived melanoma cell lines were grown in SmBM medium (Lonza, #CC-3181) supplemented with SmGM SingleQuots Supplement (Lonza, #CC4149) from the single-cell suspension of patient specimens dissociated by mechanical mincing followed by cell strainer filtering (40 mm, Thermo Scientific, #22-363-547). The established cell lines (including 10049, RRID: CVCL_F1YF; 10092, RRID: CVCL_F1YG; 10101, RRID: CVCL_E3EB; 10164, RRID: CVCL_F1YH; 10170, RRID: CVCL_F1YI; 10214, RRID: CVCL_F1YJ; 10222, RRID: CVCL_F1YK; 10266, RRID: CVCL_F1YL; 10378, RRID: CVCL_F1YM; 10417, RRID: CVCL_F1YN) underwent SNaPshot sequencing to confirm the stereotypical melanoma mutations. Murine B16F10-OVA and B16F10-*B2m* KO cell lines were kindly provided by Dr. Debattama R. Sen (MGH Cancer Center) and were cultured in DMEM-10 (Corning, #10-013-CV) supplemented with 10% FBS and 1% penicillin-streptomycin. All cell lines were authenticated and tested negative for mycoplasma and interspecies contamination using the Mycoplasma detection kit (Lonza, #LT07-318).

### TIL isolation and expansion

Patient specimens were dissociated by mechanical mincing followed by cell strainer filtering (40 µm, Thermo Scientific, #22-363-547) to generate single-cell suspension. CD8^+^ TIL were then sorted by BD FACSAria II gated on CD3^+^CD8^+^ live cells or isolated by CD8 magnetic beads (Miltenyi Biotec, #130-045-201, RRID: AB_2889920) after dead cell removal (Miltenyi Biotec, #130-090-101) or Lympholyte separation (Cedarlane Labs, #CL5020). The enriched TIL were cultured in serum-free ImmunoCult™-XF T Cell Expansion Medium (STEMCELL Technologies, #10981) with 50 ng/mL IL-2 (PeproTech, #200-02) or reactivated on plated coated with αCD3 (1 µg/mL, Bio X cell, #BE0001-2, RRID: AB_1107632) and αCD28 (1 µg/mL, Bio X cell, # BE0248, RRID:AB_2687729) for two days followed by transferring to a new plate. T cell media with IL-2 was refreshed every 2-3 days. Additionally, two TIL (10378P and 10417P) were manufactured by Iovance and banked from the leftover after the patients received the products. For lifileucel and matched baseline tumor digest single cell datasets, cells were obtained from cryopreservation, thawed, and washed twice with PBS + 0.04% BSA. Cell counts and viability were assessed and dead cell removal was performed on any sample containing fewer than 75% viable cells, using the Miltenyi Dead Cell Removal kit, according to manufacturer guidelines. Baseline tumor digests were generated via fragmentation then digestion using the Human Tumor Dissociation kit (Miltenyi PN 130-095-929) and the Miltenyi gentleMACS™ Octo Dissociator instrument according to manufacturer’s protocols. Digested cell suspensions were filtered through 70 µm cell strainers and washed using RPMI medium. CD45^+^ cells were isolated from the baseline tumor digest using the StemCell Technologies EasySep™ Release Human CD45 Positive Selection kit (PN 100-0105) and then cryopreserved for downstream analysis.

### OT-1 T cell isolation

CD8^+^ OT-I T cells were isolated from the spleens of OT-I T cell receptor transgenic mice using a CD8a T cell isolation kit (Miltenyi, #130-104-075), as per the manufacturer’s instructions. Purified OT-I T cells were then stimulated in 24-well plates coated with anti-mouse CD3ε (1 μg/mL, BioXcell, #BE0002, RRID: AB_1107630) and anti-CD28 (1 μg/mL, BioXcell, #BE0015-1, RRID: AB_1107624). After 48 hours, activated OT-I T cells were transferred into fresh media (TEXMACS medium, Miltenyi Biotec, #130-097-196) containing recombinant murine IL-2 (PeproTech, #200-02) and allowed to expand for 5-7 days.

### Flow cytometry and sorting

For flow cytometry and cell sorting, human or mouse TrueStain FcX (BioLegend, #422302 or #101320) was used to block Fc receptors before labelling cells. To discriminate live from dead cells, we used Zombie Violet Dye (BioLegend, #423114) for 20 minutes on ice, followed by surface labelling of cells for 30 minutes on ice, using standard protocols. Flow cytometry data was collected using BD FACSAria II or Cytek Northern Light. Cell sorting was performed on a BD FACSAria II instrument after staining. Analysis was performed using FlowJo software (v.10.4.1, TreeStar) using single-color compensation controls and fluorescence-minus-one thresholds to set gate margins.

### *In vitro* CRISPR screen and analysis

10049 cancer cells expressing Cas9 were transduced with Brunello whole-genome (human) lentiviral sgRNA library(58) (76,441 sgRNA, 19,114 genes, 4 guides per coding gene, 1000 control sgRNAs) at about 1000× coverage per sgRNA at MOI of 0.4. Cells were selected *in vitro* for 8 days to allow sufficient time for gene editing. A proportion of cells were then saved as Input served as a cell growth dependency control. In parallel, matched TIL were added to pre-seeded cancer cells at the E:T ratios of 0:1 (Output) or 3:1 (TIL) for 2 days. After the screen, cell pellets were lysed in ATL buffer (QIAGEN) with proteinase K (QIAGEN) before genomic DNA extraction (QIAGEN Blood Maxi kit). DNA (120 μg per sample condition to ensure 500 sgRNA representation) was PCR-amplified and sequenced using the Illumina HiSeq system. Guide sequences were demultiplexed and quantified using PoolQ v3.4.3. Read counts were normalized per 1 million reads and log2 transformed with a pseudocount of one. Gene-targeting guides were z normalized by the control sgRNA distribution(59). Guide fold changes for the 4 top performing guides were calculated as residuals fit to a natural cubic spline with 4 d.f(60). As previously described, significant depleted or enriched sgRNAs were identified using the STARS algorithm(59), considering the 4 top performing guides mapped per gene **(Supplementary Table S2)**. The CP0041 GRCh38 NCBI CRISPRko strict match chip file (12-01-2022) was used to map sgRNA barcodes to genes. Average *p*-values for genes in the library were calculated with the Hypergeometric Distribution; a control distribution for this calculation was created by grouping together 4 random control guides into pseudogenes. STRING(61) (version 12.0.) network analysis was performed on genes that scored with an FDR of < 0.05 in the TIL versus Output comparison. Clusters were determined by Markov clustering (MCL) using an inflation parameter of 3.0.

### *In vitro* cytokine treatments and growth inhibition assays

Parental and/or CRISPR-edited cancer cells were plated in flat-bottom 96-well plates (2000-4000 cells/well). The following day, cells were treated with indicated cytokines or together with IncuCyte Cytotox Green (Essen BioScience, #4633). The viable cells were enumerated after 48 hours using CellTiter-Glo (Promega, #G7570) or the dead cells were monitored by Incucyte SX5 over a course of 72 hours. For relative cell death analysis, the signal intensity from Cytotox Green was captured and normalized from control groups. Recombinant cytokines and the concentrations used in this study are listed below. Human LTα1β2 (100 ng/mL, R&D Systems, #8884-LY-025/CF), human LIGHT (100 ng/mL, BioLegend, #AF664-SP), human IFNγ (40 ng/mL, R&D Systems, #285-IF-100), human TNFα (160 ng/mL, R&D Systems, #10291-TA-100), mouse IFNγ (40 ng/mL, R&D Systems, #485-MI-100). For inhibitor studies, cells were pretreated for 2 hours with the indicated doses of ruxolitinib (MedChemExpress, #HY-50856), Nec-1s (MedChemExpress, #HY-15760), Q-VD-OPh (MedChemExpress, #HY-12305), zIETD-fmk (R&D Systems, #FMK007), Ferrostatin-1 (Sigma-Aldrich, #SML0583), Dimethyl Fumarate (Selleck Chemicals, # S2586), Granzyme B inhibitor II (Sigma-Aldrich, #368055), Perforin inhibitor (also called Perforin_IN-2, MedChemExpress, #HY-160214). All the compounds were dissolved in DMSO (0.1% final concentration). The viable cells were enumerated after 48 hours using CellTiter-Glo and were read on a Cytation 5 plate reader and analyzed by GraphPad Prism 10. All the conditions were tested in at least triplicate. The values represent the average of three to five replicates and a representative experiment from at least two independent experiments.

### *In vitro* cell cytotoxicity

For patient-derived TIL and cancer cell coculture experiments, we seed 2000-4000 cancer cells/well in 96-well plates. The next day, TIL in culture were washed with PBS and added to the cancer cells at the indicated effector: target ratios without IL-2. After 2-3 days, we washed out TIL with PBS, and the rest of the viable cancer cells were subjected to CellTiter-Glo (Promega, #G7570) analysis. For GFP or DsRed labels cancer cells cocultured with TIL, coculture experiments were set up as stated above with or without the treatment of cytokines or inhibitors as indicated in the corresponding plots. Then, the fluorescence intensity was collected in an Incucyte. For killing assays related to CD8^+^ OT-1 and B16-OVA and B16 *B2m* KO were seeded in 6-well plates pretreated with IFNγ (100 ng/mL, R&D systems) to upregulate H-2Kb. The following day, cells were trypsinized from wells, washed with PBS. B16-OVA and B16 *B2m* KO pretreated with IFN-γ were then seeded into 96-well flat-bottomed plates with 2000 cells/well. The next day, CD8^+^ OT-1 cells were washed with PBS and added to the cancer cells at the indicated effector: target ratios without IL-2. Cell viability of B16-OVA or B16 *B2m* KO was measured by CellTiter-Glo after 2-3 days. Plates were read on a Cytation 5 plate reader and analyzed by GraphPad Prism 10. All the conditions were tested in at least triplicate. The values represent the average of three to five replicates and a representative experiment from at least two independent experiments.

### Competitive cell killing

For the competitive coculture of cancer cells with TIL assays, a total of (1–2)×10^5^ cancer cells expressing GFP were seeded with a different type of cancer cells expressing DsRed at the ratio of 1:1 on the bottom of 96-well plates. On the following day, TIL in culture were washed with PBS and added to cancer cells at the indicated effector: target ratios without IL- 2. Then, the plates were placed in an Incucyte SX5 and the fluorescence intensity was collected every 3-4 hours over the time course indicated. Each condition was tested in triplicate, with 4–5 image fields analyzed per replicate. Values represent the mean of 3–5 replicates from a representative of at least two independent experiments.

### *In vitro* transwell cytotoxicity assays

*In vitro* transwell cytotoxicity experiments were performed using 0.4 μm 24-well transwell inserts (Sterlitech Corporation; # 9320412). Briefly, a total of 1 × 10^5^ cancer cells was seeded on the bottom of 24-well plates. Human CD8^+^ TIL (E:T = 4:1) or indicated control cytokines were then plated in the top compartment of the Transwell. After 48 hours of coculturing, cancer cells were washed with PBS, and the rest of the viable cancer cells were subjected to celltiterGlo analysis and were read on a Cytation 5 plate reader and analyzed by GraphPad Prism 10. All the conditions were tested in at least triplicate. The values represent the average of three to five replicates and a representative experiment from at least two independent experiments.

### Secreted cytokine profiling

Multiplexed analysis of secreted cytokines was performed using the MILLIPLEX MAP Human Cytokine/Chemokine Magnetic Bead Panel (Sigma Millipore, HCYTMAG-60K-PX30). Conditioned medium samples (25 μl) after the coculture of cancer cells and TIL were assayed. Concentration levels (pg/mL) of each protein were derived from five-parameter curve-fitting models. Fold changes relative to the control samples were calculated and plotted as the log2-transformed fold change. Lower and upper limits of quantification (LLOQ/ULOQ) were imputed from standard curves for cytokines above or below detection.

### Generation of CRISPR KO cell lines

To generate human NC, β2M KO, JAK1 KO, JAK2 KO, LTβR KO, FADDKO, CASP8 KO, TCR KO, LTβR/JAK1 DKO, β2M/LTβR/JAK1 TKO and mouse NC, TCR KO and LTβR KO cells, ribonucleoprotein (RNP)-based complexes gene editing technology was used as previously described(62). Briefly, cells were collected, pelleted, and resuspended in Opti-MEM with recombinant CAS9 protein (ThermoScientific, A36498) and synthetic guide RNA(s) (sgRNAs). Resuspended cells were electroporated at 0.5 million cells per cuvette using the Lonza 4D Nucleofector^TM^ unit with 16-well Nucleocuvette^TM^ Strips.

Gene name, sgRNA number, and sequence used in this study were as follows: *B2M, hB2M* sgRNA1, AGUCACAUGGUUCACACGGC; *B2M, hB2M* sgRNA2, UCACGUCAUCCAGCAGAGAA; *JAK1*, *hJAK1* sgRNA1, UGGUUUCAUUCGAAUGACGG; *JAK1*, *hJAK1* sgRNA2, CCGGAAGUAGCCAUCUACCA; *JAK2*, *hJAK2* sgRNA1, AGAAAACGAUCAAACCCCAC; *JAK2*, *hJAK2* sgRNA2, AAUGAAGAGUACAACCUCAG; *LTBR*, *hLTBR* sgRNA1, CUCUGCAGGUGUGAGAACCA; *LTBR, hLTBR* sgRNA2, CUCGCAGUGUGUACACUCGA; *CASP8, hCASP8* sgRNA1, GCCUGGACUACAUUCCGCAA; *CASP8*, *hCASP8* sgRNA2, UGCUUUUCCACAUCAGUCGG; *FADD, hFADD* sgRNA1, GAGGCAUAGGAACUUGAGCU; *FADD, hFADD* sgRNA2, UGACGUUAAAUGCUGCACAC; *TRAC, hTRAC* sgRNA1, AGAGUCUCUCAGCUGGUACA; *TRBC, hTRBC* sgRNA1, GCAGUAUCUGGAGUCAUUGA; control sgRNA: AUUGUUCGACCGUCUACGGG.

### Western blotting

Whole-cell lysates were lysed by RIPA Lysis Buffer (Millipore Sigma, 20-188), followed by centrifugation. Protein lysates (30-50 mg) boiled at 95 °C in 4 × fluorescence-compatible sample buffer (Invitrogen) were then loaded onto 4-12% Bolt Bis-Tris Plus gels (Life Technologies) in MES buffer (Life Technologies). Protein was transferred to a PVDF membrane using the iBLOT2 dry transferring system (Invitrogen). Membranes were blocked in Tris-buffered saline plus 0.1% Tween-20 (TBS-T) containing FL fluorescence blocking buffer (Thermo Fisher Scientific) for 1 hour at room temperature followed by overnight incubation with primary antibodies at 4°C. After washing, membranes were incubated with blocking buffer, and IRDye 800CW- or 680RD-conjugated secondary antibodies. Membranes were then visualized using the Odyssey CLx scanner (LI-COR), then analyzed using ImageJ software.

### Antibodies

For western blotting, primary antibodies against JAK1 (Cell Signaling technology, #3332, RRID:AB_2128499 and #3344 (clone 6G4), RRID:AB_2265054), JAK2 (Cell Signaling technology, #3230, RRID:AB_2128522), FADD (Cell Signaling technology, #2782S, RRID:AB_2100484), CASP8 (Cell Signaling technology, # 4790, RRID:AB_10545768), LTβR (Cell Signaling technology, #57560), and β-actin (Thermo Scientific, #MA5-15739-D800, RRID:AB_2537666 or MA5-15739-D680, RRID:AB_2537665) were used. Primary antibodies were used at 1:1,000 dilution in LI-COR Blocking Buffer. IRDye secondary antibodies against rabbit IgG, mouse IgG, or goat IgG were purchased from LI-COR Biosciences (Invitrogen) and used at 1:10,000. β-Actin was used at 1:2000 dilution as a loading control.

For *in vitro* human TIL and mouse CD8^+^ OT-1 T cells stimulation and activation, anti-human CD3 (clone OKT-3, #BE0001-2, RRID:AB_1107632, BioXcell), anti-human CD28 (clone 9.3, #BE0248, RRID:AB_2687729, BioXcell), anti-mouse CD3 (clone 17A2, #BE0002, RRID:AB_1107630, Bio X cell), and anti-mouse CD28 (clone 37.51, #BE0015-1, RRID:AB_1107624, Bio X cell) were used. For *in vitro* neutralization assay in human TIL, anti-human MHC-I (20 μg/mL, clone W6/32, RRID:AB_2561492, BioLegend), anti-human NKG2D (20 μg/mL, clone 1D11, BioXcell, RRID:AB_2894770), anti-human MICA/B (20 μg/mL, clone 6D4, RRID:AB_2715910, BioLegend), anti-human FAS (25 μg/mL, clone DX2, RRID:AB_2858235, BioLegend), anti-human IFNγ (10 μg/mL, clone B133.5, RRID:AB_2687717, BioXcell), and anti-LTα (pateclizumab, 10 μg/mL, # HY-P990034, RRID:AB_3694264, MedChemExpress) were used. For flow cytometry, the anti-human fluorochrome-conjugated antibodies used for cell surface labeling were: HLA-A, B, C (clone W6/32, RRID:AB_314878 and RRID:AB_314873, BioLegend), TCRɑ/β (clone IP26, RRID:AB_314645, BioLegend), CD3 (clone OKT3, RRID:AB_571907, BioLegend), CD4 (clone OKT4, RRID:AB_11204077, BioLegend), CD8 (clone RPA-T8, RRID:AB_2563264, BioLegend), CD39 (clone A1, RRID:AB_940427, BioLegend), TIM-3 (clone F38-2E2, RRID:AB_2561720, BioLegend), PD-1 (clone 29F.1A12, RRID:AB_2715761, BioLegend), MICA/B (clone 6D4, RRID:AB_2715910, BioLegend), FAS (CD95) (clone DX2, RRID:AB_314550, BioLegend), LTBR (clone 31G4D8, RRID:AB_2139070, BioLegend), NCR3 (Clone P30-15, RRID:AB_2149449, BioLegend). The anti-mouse fluorochrome-conjugated antibodies used for cell surface labeling were: H-2Kb (clone AF6-88.5, RRID: AB_10568693, BioLegend), TCR vβ5.1, 5.2 (clone MR9-4, RRID: AB_10897800, BioLegend).

### Multiplex IF and RNA-ISH combined with IF

*<u>Tissue Staining:</u>* FFPE tissue sections from 2 melanoma cases (10049 and 10222) were stained using a Leica BOND RX automated stainer. For each case, one section was stained with an antibody-only MIF panel and a serial section was stained with an RNA-ISH and IF combined panel. The developed MIF panel closely followed a panel previously optimised to label the following antibodies using the Opal 7-color Automation IHC Kit (NEL821001KT, Akoya Biosciences): TCF1/7 (Cell Signaling, #2203) with Opal 690, CD8 (Leica Biosystems NCL-L-CD8-4B11) with Opal 480, PD1 (Abcam ab137132) with Opal 620, PD-L1 (Cell Signaling #13684) with Opal 570, and Ki67 (Agilent M724001-2) with Opal 5201. A pool of Sox10 (Abcam ab180862, 100X dilution) and S100 (Abcam ab4066, 100X dilution) was added with detection using Opal 780 (50X dilution). In addition to the pooled SOX10/S100 antibody staining, the ISH-IF panel employed the following four RNAscope ISH probes (Advanced Cell Diagnostics): Hs-TNFSF14, Hs-LTBR-C2, Hs-LTB-C3, Hs-CD8a-C4. Corresponding Opal fluorophore concentrations (Opal 620, Opal 480, Opal 690, and Opal 520, respectively) were optimised to balance signal intensity across all channels. DAPI was used as a nuclear counterstain. *<u>Image Acquisition:</u>* Whole slide images of fluorescently stained slides were acquired on an Akoya PhenoImager HT multi-spectral slide scanner at 20x (0.496 µm/pixel) resolution. Imaging exposures were optimised for each slide to avoid pixel saturation or underexposure. Akoya inForm software was used to spectrally separate signals from each fluorophore and to mitigate the effect of native tissue autofluorescence. The spectral library used to unmix the raw images was created using the synthetic Opal spectra and autofluorescence spectra from unstained normal tonsil tissue. Unmixed tiles produced by inForm were stitched to reproduce a whole-slide pyramidal TIF using Indica Labs HALO software, which was also used to examine the multi-channel image alongside corresponding pathology-annotated H&E stained slides. *<u>Image Analysis:</u>* Quantitative image analysis was performed using the HALO image analysis platform (Indica Labs). tumor regions were verified by a pathologist and manually annotated to exclude large folds, debris, and normal tissue from quantification. The MIF and ISH-IF serial section images were co-registered and overlaid into a single image using HALO software. Quality of registration was manually verified to be 50 µm or less discrepancy between the same tissue structures. Nuclear detection and cell segmentation were performed using the HALO AI nuclear segmentation algorithm. Cell phenotyping was based on staining intensity using HALO modules HighPlex FL v4.1.3 and FISH-IF v2.2.5 for the antibody-only MIF panel and the RNA-ISH and IF combined panel, respectively. The autofluorescence (AF) channel was employed as an exclusion marker to reduce false positives from folds, debris and necrotic regions. Analysis settings were manually validated for each sample for quality of cell segmentation and phenotyping. Summary tables and object tables containing individual cell phenotype and intensity information were exported for statistical analysis **(Supplementary Table S4)**. To compare the spatial distribution of cells in 10049 and 10222, a spatial proximity analysis was performed to identify *LTBR*^+^ tumor cells (Sox10/S100^+^ *LTBR*^+^*CD8a*^-^*AF*^-^) within a 100 microns radius of each *LTB*^+^ cytotoxic T-cells (*CD8a*^+^*LTB*^+^). The average distance and total number of cells that fell within this range was then calculated. *LTBR*^+^ tumor cells were also enumerated within bands (i.e. bins) 0-50 µm micron and 50-100 µm from each *LTB*^+^ T-cell to gain better insight into potential spatial interactions **(Supplementary Table S4)**.

### Hematoxylin & Eosin Staining

#### Tissue Staining

FFPE tissue sections from 2 melanoma cases (10049 and 10222) were stained with hematoxylin and eosin (H&E) at Dana Farber/Harvard Cancer Center Specialized Histopathology Services.

#### Image Acquisition

Whole slide images of H&E-stained slides were acquired at 20x (0.543 µm/pixel) resolution using a MoticEasyScan Infinity digital pathology scanner.

### scRNA sequencing and analysis

#### Sequencing

After debris and dead cell removal, cells were counted and loaded onto the Chromium Controller (10X Genomics). Single-cell RNA libraries were generated using the 10x Genomics Chromium Single Cell V(D)J Reagent Kit using 5’ v1 chemistry with Feature Barcode technology for Cell Surface Protein (10x Genomics; Cat# 1000080). After each step, cDNA generation, gene expression libraries, and cell surface protein libraries samples quality was assessed using the Qubit dsDNA high sensitivity kit (Invitrogen; Cat# Q32854) and the high sensitivity BioA DNA kit (Agilent; Cat# 5067-4626). Samples that passed quality control were sequenced on Novaseq 6000 XP (Illumina), using pair-end reads, with 26 reads for read 1 and 90 reads for read 2, or were sequenced on a NextSeq 500 sequencer (Illumina), using pair-end reads, with 26 reads for read 1 and 55 reads for read 2. For lifileucel and matched baseline tumor digest single cell dataset, sequencing libraries were prepared using 10x 5’ Immune profiling v2 reagents, following manufacturers guidelines; and sequenced on Illumina NextSeq2000 instrument with an insert read length of 90 base pairs.

#### scRNA-seq read alignment and quantification

Raw sequencing data for expanded TILs and post-coculture samples were pre-processed using Cell Ranger Multi (v9.0.1, 10x Genomics) to analyze 5′ single-cell RNA-seq and antibody-derived tag (ADT) libraries from fifteen multiplexed samples across three pools. Reads were aligned to the GRCh38 reference genome (v3.0.0, 10x Genomics), and unique molecular identifiers (UMIs) were counted to generate a cell-by-gene count matrix(63). Sample demultiplexing was performed using hashtag oligonucleotides (HTOs) from the TotalSeq-C™ panel (BioLegend). For the paired S_10049 TIL and tumor samples and the OT-1 mouse dataset, pre-processing was performed using Cell Ranger (v7.0.0, 10x Genomics). FASTQ reads were demultiplexed, aligned to either the GRCh38 (human) or mm10 (mouse) reference genomes, and used to generate UMI count matrices. Low-quality droplets were filtered out of the matrix prior to proceeding with downstream analyzes using the percent of mitochondrial UMIs and number of unique genes detected as filters (S_10049 paired TIL and tumor samples: <15% mitochondrial UMIs, > 200 unique genes; expanded TILs and cocultured TILs: <15% mitochondrial UMIs, > 400 unique genes; OT-1 mouse samples: <10% mitochondrial UMIs, >400 unique genes; lifileucel and matched baseline tumor digest single cell samples: <10% mitochondrial UMIs, > 200 and < 6000 unique genes.). The percent of mitochondrial UMI was computed using 13 mitochondrial genes for human data (MT-ND6, MT-CO2, MT-CYB, MT-ND2, MT-ND5, MT-CO1, MT-ND3, MT-ND4, MT-ND1, MT-ATP6, MT-CO3, MT-ND4L, MT-ATP8) and 35 mitochondrial genes for mouse data (mt-Tf, mt-Tv, mt-Ti, mt-Tq, mt-Tm, mt-Tw, mt-Ta, mt-Tn, mt-Tc, mt-Td, mt-Tk, mt-Tg, mt-Tr, mt-Th, mt-Te, mt-Tt, mt-Tp, mt-Ts1, mt-Rnr1, mt-Rnr2, mt-Tl1, mt-Nd13, mt-Nd4, mt-Nd5, mt-Nd6, mt-Nd4l, mt-Cytb, mt-Tl2, mt-Nd2, mt-Nd, mt-Nd2, mt-Co1, mt-Co2, mt-Co3, mt-Atp6, mt-Atp8) using the qc_metrics function in Pegasus(64) for expanded TILs, matched S_10049 TIL and tumor, and OT-1 datasets, and the PercentageFeatureSet function in Seurat v5(65) for the lifileucel and matched baseline tumor digest datasets. The counts for each remaining cell in the matrix were then log-normalized by computing the log1p (counts per 100,000), which we refer to in the text and figures as logCPM. *<u>Basic clustering</u>*: For expanded TIL, paired S_10049 TIL and tumor, and OT-1 mouse data, 2,000 highly variable genes were selected using the *highly variable features* function in Pegasus and used as input for principal component analysis. The resulting principal component scores were aligned using the Harmony algorithm to account for technical variability between donors(66). The resulting principal components were used as input for Leiden clustering and Uniform Manifold Approximation and Projection (UMAP) algorithm (spread = 1, min-dist = 0.5). The number of principal components used for each S_10049 TIL clustering (6), S_10049 tumor clustering (28), and mouse OT-I data (14) was decided via molecular cross-validation(67). For lifileucel and matched baseline tumor digest single cell datasets, the default Seurat clustering and visualization pipeline was used with the following parameter modifications: number of PCs = 75, integration method = Harmony, clustering resolution = 0.5 (lifileucel) or 1 (baseline tumor digest).

#### Atlas of melanoma TIL

Publicly available melanoma TIL datasets were downloaded and integrated with the expanded TIL dataset (*n* = 5) to further characterize the immune landscape. Cells from Chiffelle et al.(68), 2024, Oliveira et al., 2021(50),Jerby-Arnon et al., 2018(49) and baseline tumor digest from S_10049 underwent quality control with metrics <15% mitochondrial UMIs, >400 unique genes for 10X datasets, and author provided post-QC assignments for SmartSeq2 datasets.. The top 1000 highly variable genes were selected using Scanpy’s highly_variable_genes() method with seurat_v3 flavor, accounting for inter-dataset batch effects, and intra-dataset batch effects when applicable. We then used scVI to learn a 30-dimensional batch-corrected latent representation, specifying individual samples as a discrete covariate. The model was trained for 300 epochs with two hidden layers. The learned latent space was used to compute a neighborhood graph, followed by Uniform Manifold Approximation and Projection (UMAP) algorithm (spread = 1, min-dist = 0.5) for visualization and Leiden clustering to identify transcriptionally distinct cell populations. CD8+ T cells were identified from this embedding by examining the expression of *CD3E*, *CD3D*, *CD8A*, *CD8B, NCAM1* and *CD4*, and the above steps repeated to generate the embedding and clustering.

#### Cell-cell interaction analysis

To infer potential receptor: ligand interactions between cell-cell pairs, we used CellPhoneDB(47) (v2.1.7, Python) and ran the algorithm on an aggregated counts matrix of TIL and tumor cells. Both TIL and tumor were tested as both a sender (ligand) and receiver (receptor) populations as defined by the algorithm. We restricted potential interactions to those where the receptor and ligand were each expressed in >20% of their respective sample, with significance defined as an empirical *P* < 0.001 cutoff. Mean was defined by the algorithm as the aggregate mean expression (logCPM) of the receptor and ligand genes in the cell-cell pair. A curated list of interactions between TIL and tumor were visualized as dot plots using the ggplot2 package (v3.3.3, R). The full list of predicted receptor-ligand interactions is compiled in ***Supplementary Table S3***.

#### Marker gene identification and cell annotation

The marker genes defining each distinct cell subset from our scRNAseq datasets were determined using two complementary methods. First, we calculated the area under the receiver operating characteristic (AUROC) curve for the logCPM values of each gene as a predictor of subset membership using the *de_analysis* function in Pegasus. Genes with an AUROC ≥ 0.75 were considered marker genes for a particular cell subset. Second, we created a pseudobulk count matrix(69) by summing the UMI counts across cells for each unique cluster/sample combination, creating a matrix of n genes x (n samples*n clusters). We performed “one-versus-all” (OVA) differential expression (DE) analysed for each cell subset using the Limma package (v3.54.0, R)(70). For each subset, we used an input model *gene ∼ in_clust*, where *in_clust* is a factor with two levels indicating if the sample was in or not in the subset being tested. A moderated t-test was used to calculate p values and compute a false discovery rate (FDR) using the Bejamini-Hochberg method. We identified marker genes that were significantly associated with a particular subset as having an FDR < 0.05 and a log2 fold change > 0. For lifileucel and matched baseline tumor digest single cell datasets, cluster marker genes were identified using the FindAllMarkers function in Seurat with default parameters. Marker genes for each cell subset were interrogated and investigated in the context of other published immune profiles to guide our cell subset annotations. Thresholds used to define cell subsets as presented in Figure 4 are as follows: LTB^+^CD8^+^ T cell is single cell expression greater than or equal to 1 normalized count for *LTB* while Baseline_LTB^hi^ and Baseline_NeoTCR8^hi^ are greater than or equal to the 90th percentile of expression for *LTB* and the NeoTCR8 gene signature respectively.

### TCR repertoire reconstruction and analysis

For expanded TIL dataset, Immune repertoire reconstruction was done using open-source TRUST4 v.1.0.8(71)(https://github.com/liulab-dfci/TRUST4) with default arguments. Cells with TCRβ chain sequences with full V, J, and CDR3 information were retained for downstream analysis. Expanded clones were defined as cells with a unique TCRβ chain that was recorded in >10% of cells with TRUST4 reconstructed TCR repertoires and recovered on > 5 cells per patient at a given timepoint. The number of TCR clones and recovered V, J, and CDR3 information can be found in ***Supplementary Table S6***.

### Differential gene expression analysis

Differential gene expression analysis was performed on pseudobulk count matrices using the DESeq2 package (v. 1.38.2, R) or Limma package (v3.54.0, R)(70). The input model was *gene ∼ comparison,* where *comparison* was a binary variable. Differentially expressed genes were selected based on a cutoff of Log2FoldChange > 1 or < −1 and FDR < 0.1. All differential gene expression analysis results can be found in ***Supplementary Table S5 and S7***.

### Gene set scoring analysis

For lifileucel gene set scoring, patient-cluster-level pseudobulk profiles were generated using the Aggregate Expression function in Seurat then rank-based single-sample gene set scores quantified for the CD39 CD69 double negative (stem-like) and double positive (terminal differentiated) gene sets from Krishna et al.(6) using the R package singscore(72,73) with default parameters. For matched baseline tumor digest single cell gene set scoring, the R packages Escape(74) and UCell(75) were used to quantify the gene set scores for the NeoTCR8 gene signature from Lowery et al.(54) with default parameters.

### Statistical Methods, Data Analysis, and Software

All graphs with error bars (shaded region) report mean ± s.d. values except where indicated. Statistical tests and number of replicates are listed in the text and figure legends. All the key conclusions have been confirmed in 2-3 independent experiments with multiple (n > 3) biological replicates. Multiple-hypothesis testing correction was applied where multiple hypotheses were tested and is indicated by the use of FDR. The data analysis software used included FlowJo (v10, RRID:SCR_008520), Graphpad Prism (v10, RRID:SCR_002798), Image J (v.2.9.2, RRID:SCR_003070), Cell Ranger (v7.0.0 and v.9.0.1, RRID:SCR_017344), CellPhoneDB (v4.1.0, RRID:SCR_017054), anndata (v.7.5.6, R, RRID:SCR_018209), ggplot2 (v3.5.1, R, RRID:SCR_014601), Limma (v3.54.2, RRID:SCR_010943), TRUST4 (v.1.0.8, RRID:SCR_026162), DEseq2 (v.1.38.2, RRID:SCR_015687), pegasus (1.10.2, Python, RRID:SCR_021645), scanpy (1.10.3, Python, RRID:SCR_018139), matplotlib (3.9.4, Python, RRID:SCR_008624), pandas (2.3.0, Python, RRID:SCR_018214), numpy (2.0.2, Python, RRID:SCR_008633), scipy (1.13.1, Python, RRID:SCR_008058), anndata (0.10.9, Python, RRID:SCR_018209), PoolQ (v3.4.3), STRING (v12.0, RRID:SCR_005223).

## Supporting information

Supplementary Figures with Legends

## Data availability

The datasets generated and analyzed in this study are included in the article and its Supplementary Information. scRNA-seq datasets of human TIL have been deposited in GEO under the accession GSE26883, except a sub-data set on Lifileucel and matched tumor that have not been made available publicly in a repository to protect the privacy and confidentiality of the participating patients. Upon reasonable request and authorization by Iovance Biotherapeutics, and after appropriate data-sharing and use agreements have been agreed upon, eligible academic researchers in the field may be provided access to de-identified gene-limited datasets. Requesters should submit a proposal outlining the objective, data format and features, hypothesis and specific rationale to. Requests will usually be processed within 15 weeks.

## Code availability

The codes used to figure generation and data analysis in this study are available upon request. The code used to generate figures for all scRNA-seq data analysis are present here https://github.com/villani-lab/mhc_class_i_independant_cancer_cell_lysis and will be available upon publication. For analysis of lifileucel and matched baseline tumor digest single cell dataset, no original code was created. Publicly available packages were used and code modified from publicly available sources for case-specific use.

## Acknowledgements

This work was supported by NIH 1R37CA283560-01 (R.W.J.), and the MGH ECOR Fund for Medical Discovery (FMD) Research Fellowship Award (H.X.). Additional support provided by a generous gift from Robert and Marie McInnes. We acknowledge funding provided by the Krantz Family Center for Cancer Research Breakthrough Award (M.S-F., G.M.B, R.W.J.), the National Institute of Health Director’s New Innovator Award (DP2CA247831; A.C.V.), the Massachusetts Life Sciences Center Research Infrastructure Program in support of the Mass General Cancer Center Tumor Cartography Center, and the Dr. Miriam and Sheldon G. Adelson Medical Research Foundation. D.B.K. is supported by U54-CA272688, R01-HL157174, R01-CA285308, R01-CA279391 & R01NS140967. The funding bodies had no role in the design of the study, and collection, analysis, and interpretation of the data, or in writing the manuscript. The authors thank all members of the Sen, Villani, Manguso, and Jenkins laboratories at MGH and the Broad Institute of MIT and Harvard.

## Author Contributions

*Conception and experimental design*: H.X., R.T.M, and R.W.J.

*Methodology and data acquisition*: H.X., A.J., A.D., J.W.D., J.P., N.S., A.C., S.A., A.M.C., Y.S., W.A.M., M.F., J.F., O.Y.R., K.H.X., Y.S., B.P., J.D.S., W.J.L., D.B.K., L.T.N., M.S.F., and R.W.J.

*Analysis and interpretation of data*: H.X., A.D., J.W.D., J.P., N.S., S.A., B.P., D.B.K., L.T.N., R.Q., H.Y., A-C.V., R.T.M., and R.W.J.

*Administrative, technical, or material support*: H.X., T.S., A.L., S.C., D.P.L., R.J.S., K.T.F., G.M.B., N.H., D.R.S., C.W., B.G., R.Q., H.Y., A-C.V., R.T.M., and R.W.J.

*Manuscript writing and revision*: H.X., A.D., J.W.D., N.S., A.C., R.J.S., K.T.F., H.Y., A-C.V., R.T.M., and R.W.J.

**Correspondence and requests for materials** should be addressed to R.W.J.

## Supplementary Information Guide

### Supplementary Figure Legends

**Supplementary Fig. S1 | Patient-derived TIL immunophenotyping**. **A-K**, Immunophenotyping (CD4, CD8, CD39, TIM-3, PD-1) of expanded in-house TIL including (**A**) 10049, (**B**) 10092, (**C**) 10101, (**D**) 10164, (**E**) 10170, (**F**) 10214, (**G**) 10222, (**H**) 10266, (**I**) 10378, and TIL manufactured by Iovance including (**J**)10378P and (**K**) 10417P.

**Supplementary Fig. S2 | MHC I-independent killing of cancer cells by TIL**. **A**, UMAP embedding of scRNA-seq data for indicated TIL samples. **B**, Heatmap showing the representative marker genes that are differentially expressed across different clusters. **C**, Dot plot showing the expression of indicated marker genes across clusters. **D-H**, Cell viability of (**D**) 10101, (**E**) 10164, (**F**) 10170, (**G**) 10214, and (**H**) 10222 control (sg*NC*) and *B2M*-null (sg*B2M*) melanoma cell lines cocultured with autologous TIL at the indicated E:T ratios for 2 days. **I-K**, Cell viability of (**I**) 10049, (**J**) 10214, and (**K**) 10378 control (sg*NC*) and *B2M*-null (sg*B2M*) melanoma cell lines cocultured with autologous TIL at the indicated E:T ratios for 1, 2, and 3 days. **L**, Representative images of Fig. 2d and 2e showing the viable 10049 sg*NC*-GFP and sg*B2M*-DsRed in competitive coculture experiments as indicated. **M-O,** Representative images **(M)** and normalized fluorescence intensity of viable 10378 sg*NC*-GFP and sg*B2M*-DsRed cells from a competitive coculture with (**N**) autologous TIL or (**O**) TCR KO (sg*TRAC* and sg*TRBC*) autologous TIL at the indicated E:T ratios and over the time course of 99 hours. Panels **D-K** and **N-O** are normalized to control groups, and *P* values were tested by two-way ANOVA with Tukey correction of multiple comparisons. Mean +/− s.d. (bar or shaded region) is shown (*n* = 3). \**P* < 0.05; \*\**P* < 0.01; \*\*\**P* < 0.001; **** *P* < 0.0001; *ns*, not significant.

**Supplementary Fig. S3 | TIL-mediated killing requires cell contact, but not unconventional T cells, FAS-FASL, NKG2D-NKG2DL, or granzyme B/perforin**. **A**, Cell viability of 10049 melanoma cells treated with TIL at E:T ratios of 0:1, 4:1 or conditioned media collected after the coculture of cancer cells and TIL (E:T = 4:1) for 2 days. Individual values (open circles) are shown (*n* = 3; one-way ANOVA with Tukey correction of multiple comparisons). **B**, Dot plot showing the expression of indicated marker genes across clusters of expanded TIL (corresponding to Fig. 1D). **C**, Flow cytometry plots showing the percentage of NCR3^+^CD8^+^ T cells in 10049, 10214, 10378P, and 10417P TIL. **D**, UMAP embedding depicting the clusters of 10049 cancer cell scRNA-seq. **E**, UMAP embedding depicting the single-cell expression of *FAS* in melanoma cells. **F**, Flow cytometry histogram of FAS expression +/− treatment of FAS neutralizing antibodies in 10049 melanoma cells. **G-J**, Cell viability assessment of (**G**) 10049, (**H**) 10214, (**I**) 10378, and (**J**) 10417 control (sg*NC*) and *FAS*-null (sg*FAS*) melanoma cell lines cocultured with autologous TIL at indicated E:T ratios for 1, 2, and 3 days. **K**, UMAP embedding depicting the single-cell expression of *MICA* and *MICB* in 10049 melanoma cells. **L**, Flow cytometry histogram of MICA/B expression +/− treatment with MICA/B neutralizing antibodies in 10049 melanoma cells. **M**, Representative images of viable 10049-GFP melanoma cells +/− treatment with the Granzyme B II inhibitor (15 nM) or perforin inhibitor (Perforin_IN-2, 10 µM) cocultured with autologous TIL at the indicated E:T ratios for 2 days. In panels **G-J,** mean +/− s.d. (bar) is shown (*n* = 3; two-way ANOVA with Tukey correction of multiple comparisons).

**Supplementary Fig. S4 | LTβR and IFN sensing pathways are involved in TIL-mediated killing**. **A-B**, Library recovery. Dotted lines represent the Gaussian distribution of (**A**) sgRNA abundance and (**B**) sgRNA frequency (input, output, and TIL-treated groups are shown in solid lines. **C-D**, Effect size model for sgRNAs. A natural cubic spline (solid line) was fit between (**C**) input and output and (**D**) between TIL and output sgRNA abundances, and the effect size is calculated as the residual from the spline. **E**, A matrix of Pearson’s correlations of the library distribution from one technical replicate compared to any other technical replicate across input, output, and TIL conditions. **F**, Genes ranked by output versus input-normalized fold change for genome-scale screens, with circle size corresponding to FDR by STARS. Genes with 4-5 mapped guides are visualized; fold changes are calculated only from 4 top-performing guides per gene. Selected top genes with FDR values < = 0.3 at each end are listed and arranged by statistical significance. **G**, Frequency histograms of enrichment or depletion (*z* score) for the selected sgRNAs. sgRNAs targeting indicated genes are shown by the green lines. For genes with more than 4 mapped guides, only the top 4 performing guides were used for fold change and STARS calculations. **H**, STRING network analysis of the depleted and enriched genes with FDR < 0.05, with circles size-scaled by −log_10_(STARS score) and colored by average gene log fold-change. **I**, Volcano plot highlighting the relative depletion and enrichment of sgRNAs targeting *TNFRSF1A (TNFR1)*, *FAS*, and *TNFSF10* (TRAIL). **J-K**, Western blot for JAK1, LTβR, and β-actin in CRISPR-edited **(J)** 10049 and **(K)** 10378 melanoma cells with indicated sgRNA pairs. **L-M**, Viability assessment of CRISPR-edited (**L**) control (sg*NC*) and (**M**) sg*B2M*-null (sg*B2M*) 10378 melanoma cells with indicated sgRNA pairs cocultured with autologous CD8^+^ TIL at indicated E:T ratios for 1, 2, and 3 days compared to untreated controls. Mean +/− s.d. (bar) is shown (*n* = 3; Two-way ANOVA with Geisser-Greenhouse correction and Tukey correction for multiple comparisons). \**P* < 0.05; \*\**P* < 0.01; \*\*\**P* < 0.001; **** *P* < 0.0001; *ns*, not significant.

**Supplementary Fig. S5 | TIL-mediated killing is via cell-extrinsic apoptosis and necroptosis**. **A,** Representative images from competition assays using equal proportions of (*left*) control (GFP) and LTβR/JAK1 DKO (DsRed) or *B2M*-null (GFP) and (*right*) *B2M*-null/LTβR/JAK1 TKO (DsRed) 10049 melanoma cells co-cultured with autologous TIL at different E:T ratios at indicated times (corresponds to Fig. 3E and 3F). **B,** Representative images from competition assays using equal proportions of (*left*) control (GFP) and LTβR/JAK1 DKO (DsRed) or *B2M*-null (GFP) and (*right*) *B2M*-null/LTβR/JAK1 TKO (DsRed) 10378 melanoma cells co-cultured with autologous TIL at different E:T ratios at indicated times (corresponds to C and D). **C-D**, Competition assays using equal proportions of (**C**) control (GFP) and LTβR/JAK1 DKO (DsRed) or (**D**) *B2M*-null (GFP) and *B2M*-null/LTβR/JAK1 TKO (DsRed) 10378 melanoma cells co-cultured with autologous TIL at different E:T ratios at indicates times. Normalized fluorescence intensity of viable cancer cells (mean +/− s.d.; shaded region) is shown (*n* = 3, two-way ANOVA with Tukey correction of multiple comparisons). Inset stacked bar blot (E:T = 4:1) depicts proportional decrease in GFP and DsRed cells over time. **E**, Cell viability assessment of control (sg*NC*) and *B2M*-null (sg*B2M*) 10049 melanoma cells pretreated with/without JAK1/2 inhibitor (ruxolitinib, 1 μM) for 3 hours followed by coculturing with autologous TIL (E:T = 4:1) for 2 days compared to untreated control. Individual values (open circles) are shown (*n* = 3; 2-sided paired *t*-test). **F-G**, Western blot for (**F**) FADD and (**G**) caspase 8 in CRISPR-edited 10049 melanoma cells with 2 different sgRNA per gene target (with β-actin as loading control). **H**, Cell viability assessment of control (sg*NC*), *CASP8*-null (sg*CASP8*), and *FADD*-null (sg*FADD*) 10049 melanoma cells cocultured with autologous TIL (E:T 4:1) for 2 days compared to untreated control. **I-J**, Cell viability assessment of melanoma cells (10049) pretreated (3 hours) with Q-VD-OPh (10 μM), zIETD-fmk (10 μM), Nec-1s (10 μM), DMF (dimethyl fumarate, 25 μM), Ferrostatin-1 (2 μM), or combinations thereof as indicated prior to coculture with autologous TIL (E:T 4:1) for 2 days, compared to vehicle control (0.1% DMSO). Individual values (open circles) are shown (*n* = 3; one-way ANOVA with Tukey correction of multiple comparisons). **K-L**, Viability assessment of CRISPR-edited 10049 melanoma cells with indicated sgRNA treated with recombinant LIGHT (50 ng/mL), IFNγ (40 ng/mL) alone or in different combinations as indicated for 2 days. **M**, Western blot for JAK2 in CRISPR-edited 10049 melanoma cells with 2 different sgRNA (β-actin as loading control). **N-Q**, Viability assessment of CRISPR-edited 10049 melanoma cells with indicated sgRNA treated with recombinant IFNγ (40 ng/mL) +/− LIGHT (50 ng/mL) or LTɑ1β2 (50 ng/mL) for 2 days compared to untreated controls. Mean values (bars) and individual values (open circles) are shown (*n* = 4; 2-sided paired *t*-test). \**P* < 0.05; \*\**P* < 0.01; \*\*\**P* < 0.001; **** *P* < 0.0001; *ns*, not significant.

**Supplementary Fig. S6 | Melanoma cell lines show sensitivity to recombinant LTBR ligands and IFNγ**. **A-P**, Cell viability assessment of (**A-B**) 10049, (**C-D**) 10092, (**E-F**) 10101, (**G-H)** 10164, (**I-J**) 10170, (**K-L**) 10214, (**M-N**) 10222, and (**O-P**) 10266 melanoma cell lines treated with recombinant LIGHT (50 ng/mL) or LTɑ1β2 (50 ng/mL) +/− IFNγ (40 ng/mL) as indicated for 2 days compared to untreated controls. Mean +/− s.d. (bar) is shown (*n* = 3; Two-way ANOVA with Geisser-Greenhouse correction and Tukey correction for multiple comparisons). **Q**, Flow cytometry histogram of LTβR expression in indicated melanoma cell lines. **R-T**, UMAP embedding depicting the single-cell co-expression of (**R**) *LTB* and *LTA,* (**S**) *LTB*, *IFNG*, and *LTA*, and (**T**) *IFNG* and *TNFSF14* in expanded TIL (10049). \**P* < 0.05; \*\**P* < 0.01; \*\*\**P* < 0.001; **** *P* < 0.0001; *ns*, not significant.

**Supplementary Fig. S7 | Dynamic lymphotoxin expression during expansion and upon coculture with cancer cells**. **A**, H&E and immunofluorescence (IF) with RNA *in situ* hybridization (RNA-ISH) showing the expression of *CD8a*, *LTB*, *TNFSF14*, *LTBR*, and SOX10/S100 in 10049 tumor tissues. **B-C**, Percentage of SOX10/S100^+^ melanoma cells and *CD8a*^+^ T cells and their subpopulations in (**B**) 10049 and (**C**) 10222 specimens, respectively. **D**, Percentage of SOX10/S100^+^*LTBR*^+^ melanoma cells and *LTB*^+^*CD8*^+^ T cells in the cells analyzed in 10049 tumor specimen. **E**, Percentage of SOX10/S100^+^*LTBR*^+^ melanoma cells within the indicated distance of *LTB*^+^*CD8a*^+^ T cells in the 10049 tumor specimen. **F**, Stacked bar plot showing the relative abundance of cells from various datasets across clusters related to Fig. 4E. **G**, Heatmap showing the representative marker genes differentially expressed across clusters related to Fig. 4E. **H**, UMAP embedding depicting the samples included related to Fig. 5A. **I,** Heatmap showing the representative marker genes differentially expressed across clusters related to Fig. 5A. **J,** Abundance difference in clusters of TIL before and after coculture with sg*NC* (left) and sg*B2M* (right) melanoma cells. **K,** UMAP embedding depicting expanded TCR clones (calculated by TCR β chains). **L**, Scatter plots showing the proportion of TCR clones before and after coculture with sg*NC* (left) and sg*B2M* (right) cancer cells. **M**, Log2 fold changes of *LTA* expression across clusters before and after coculture control (sg*NC*) and *B2M*-null (sg*B2M*) melanoma cells. **N**, Bar plot showing the percentage of *LTA*^+^*CD8*^+^ cells in CD8^+^ TIL before and after coculture with cancer cells transfected with sg*B2M* (*n* = 2). **O**, Dot plot showing selected genes before and after coculture with B2M-nul (sg*B2M*) cancer cells. **P**, Bar plot examining the log2 fold change (mean CPM) of *LTA*/*LTB* ratio in TIL before and after coculture with sg*B2M*.

**Supplementary Fig. S8 | Neutralization of LTɑ and IFNγ rescue cancer cells from TIL-mediated killing. A**, Flow cytometry histogram of H-2Kb in B16-OVA or *B2m* KO cells with/without being pretreated with IFNγ (100 ng/mL, 24 hours). **B**, Scatter plot showing the differential gene expression that encodes the surface proteins in OT-1_0 versus OT-1_24 cells (*top*) and OT-1_0 versus OT-1_48 cells (*down*). **C-D,** Normalized fluorescence intensity of viable melanoma cells (10049, sg*B2M*-DsRed) cocultured with autologous TIL and treated with or without the indicated neutralizing antibodies (anti-LTα, 10 μg/mL; anti-IFNγ, 10 μg/mL) at E:T ratio of (**C**) 1:1 or (**D**) 2:1 for the indicated time points.

### Supplementary Tables -

**Supplementary Fig. S9 | The abundance of *LTB*^+^*CD8*^+^ T cells in the expanded TIL products is associated with response to TIL therapy. A**, UMAP embedding of scRNA-seq data for TIL depicting the distribution of patients who showed response (R), stable disease (SD), or progression disease (PD) after receiving TIL therapy. **B**, Dot plots showing the representative marker genes differentially expressed across clusters in baseline tumor digests.

**Supplementary Table S1-** Summary of demographic characteristics of patient specimens used for melanoma cell line generation and TIL expansion. Supporting Fig. 1 and Supplementary Fig. S1

**Supplementary Table S2-** Enriched and depleted guide RNAs calculated by STAR algorithm compared to TIL and Output, or Input and Output. Supporting Fig. 3 and Supplementary Fig. S4

**Supplementary Table S3-** Ligands and receptors analysis using paired melanoma cells and TIL. Supporting Fig. 3

**Supplementary Table S4-** Summary of tumor and T cell ratio and proximity analysis. Supporting Fig. 5 and Supplementary Fig. S7.

**Supplementary Table S5-** Differential gene analysis of CD8^+^ TIL between baseline tumor digests and expanded TIL. Supporting Fig. 5 and Supplementary Fig. S7.

**Supplementary Table S6-** Number of TCR colones before and after coculture, and recovered V, J, and CDR3 segment information. Supporting Fig. 5 and Supplementary Fig. S7

**Supplementary Table S7-** Differential gene analysis of OT-1_24 versus OT-1_0 cells, and OT-1_48 versus OT-1_0 cells. Supporting Fig. 5 and Supplementary Fig. S8

