## Supplementary Figures with Legends for "Lymphotoxin-driven cancer cell eradication by tumoricidal CD8^+^ TIL"

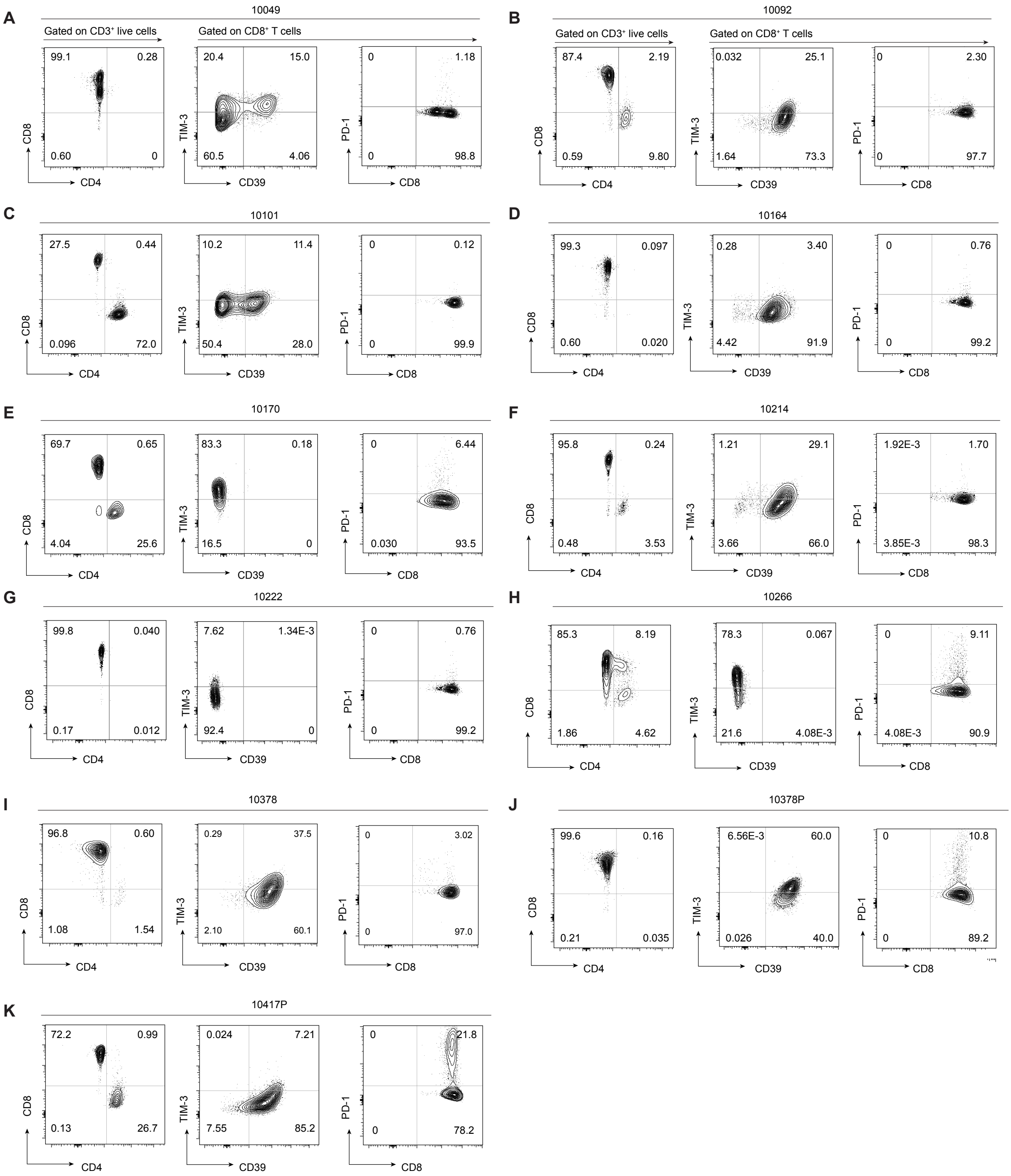


**Supplementary Fig. S1 | Patient-derived TIL immunophenotyping**. **A-K**, Immunophenotyping (CD4, CD8, CD39, TIM-3, PD-1) of expanded in-house TIL including (**A**) 10049, (**B**) 10092, (**C**) 10101, (**D**) 10164, (**E**) 10170, (**F**) 10214, (**G**) 10222, (**H**) 10266, (**I**) 10378, and TIL manufactured by Iovance including (**J**)10378P and (**K**) 10417P.

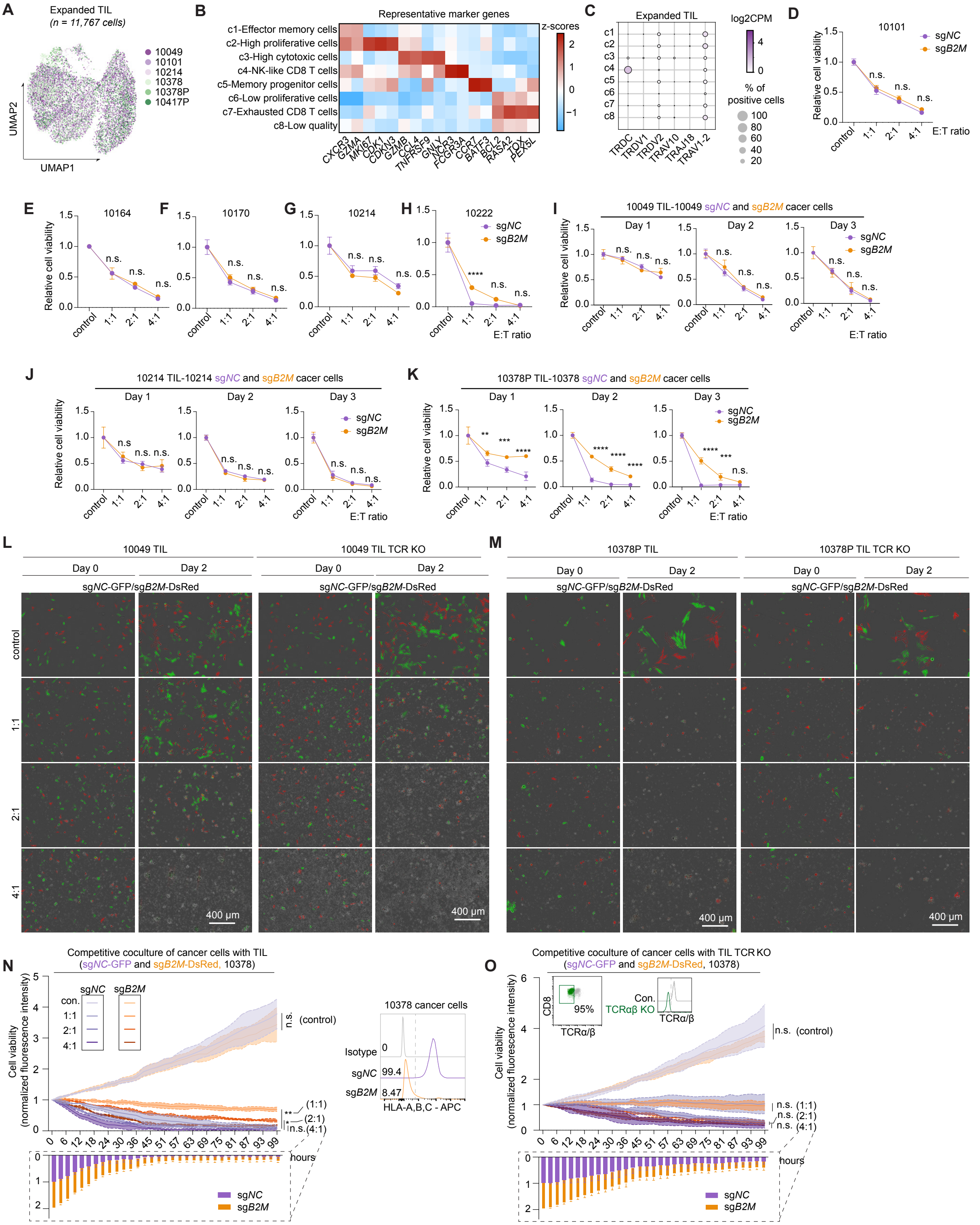


**Supplementary Fig. S2 | MHC I-independent killing of cancer cells by TIL**. **A**, UMAP embedding of scRNA-seq data for indicated TIL samples. **B**, Heatmap showing the representative marker genes that are differentially expressed across different clusters. **C**, Dot plot showing the expression of indicated marker genes across clusters. **D-H**, Cell viability of (**D**) 10101, (**E**) 10164, (**F**) 10170, (**G**) 10214, and (**H**) 10222 control (sg*NC*) and *B2M*-null (sg*B2M*) melanoma cell lines cocultured with autologous TIL at the indicated E:T ratios for 2 days. **I-K**, Cell viability of (**I**) 10049, (**J**) 10214, and (**K**) 10378 control (sg*NC*) and *B2M*-null (sg*B2M*) melanoma cell lines cocultured with autologous TIL at the indicated E:T ratios for 1, 2, and 3 days. **L**, Representative images of Fig. 2d and 2e showing the viable 10049 sg*NC*-GFP and sg*B2M*-DsRed in competitive coculture experiments as indicated. **M-O,** Representative images **(M)** and normalized fluorescence intensity of viable 10378 sg*NC*-GFP and sg*B2M*-DsRed cells from a competitive coculture with (**N**) autologous TIL or (**O**) TCR KO (sg*TRAC* and sg*TRBC*) autologous TIL at the indicated E:T ratios and over the time course of 99 hours. Panels **D-K** and **N-O** are normalized to control groups, and *P* values were tested by two-way ANOVA with Tukey correction of multiple comparisons. Mean +/- s.d. (bar or shaded region) is shown ($n = 3$). *$P < 0.05$; **$P < 0.01$; ***$P < 0.001$; **** $P < 0.0001$; *ns*, not significant.

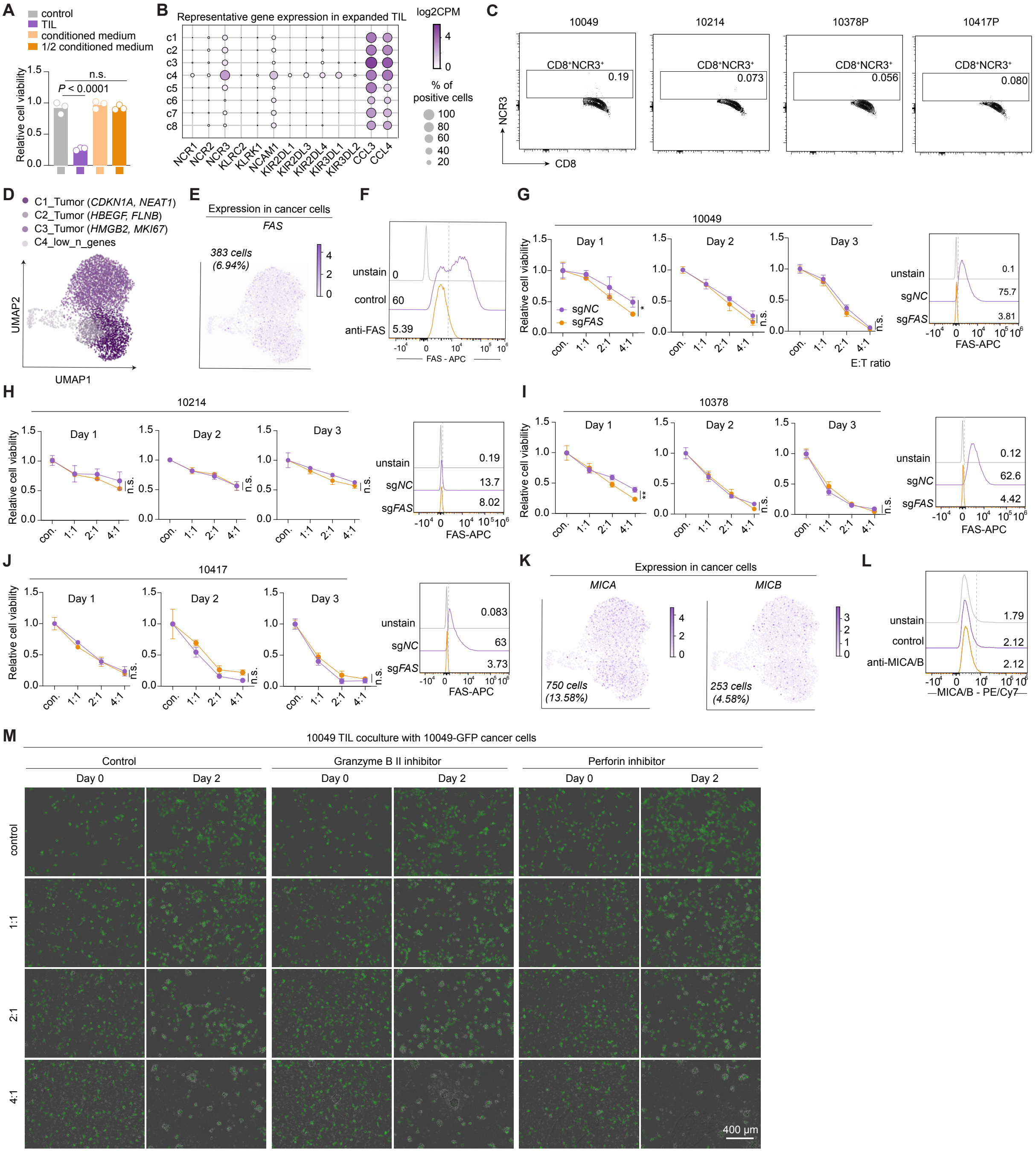

**Supplementary Fig. S3 | TIL-mediated killing requires cell contact, but not unconventional T cells, FAS-FASL, NKG2D-NKG2DL, or granzyme B/perforin**. **A**, Cell viability of 10049 melanoma cells treated with TIL at E:T ratios of 0:1, 4:1 or conditioned media collected after the coculture of cancer cells and TIL (E:T = 4:1) for 2 days. Individual values (open circles) are shown ($n$ = 3; one-way ANOVA with Tukey correction of multiple comparisons). **B**, Dot plot showing the expression of indicated marker genes across clusters of expanded TIL (corresponding to Fig. 1D). **C**, Flow cytometry plots showing the percentage of NCR3$^+$CD8$^+$ T cells in 10049, 10214, 10378P, and 10417P TIL. **D**, UMAP embedding depicting the clusters of 10049 cancer cell scRNA-seq. **E**, UMAP embedding depicting the single-cell expression of *FAS* in melanoma cells. **F**, Flow cytometry histogram of FAS expression +/- treatment of FAS neutralizing antibodies in 10049 melanoma cells. **G-J**, Cell viability assessment of (**G**) 10049, (**H**) 10214, (**I**) 10378, and (**J**) 10417 control (sg*NC*) and *FAS*-null (sg*FAS*) melanoma cell lines cocultured with autologous TIL at indicated E:T ratios for 1, 2, and 3 days. **K**, UMAP embedding depicting the single-cell expression of *MICA* and *MICB* in 10049 melanoma cells. **L**, Flow cytometry histogram of MICA/B expression +/- treatment with MICA/B neutralizing antibodies in 10049 melanoma cells. **M**, Representative images of viable 10049-GFP melanoma cells +/- treatment with the Granzyme B II inhibitor (15 nM) or perforin inhibitor (Perforin_IN-2, 10 μM) cocultured with autologous TIL at the indicated E:T ratios for 2 days. In panels **G-J,** mean +/- s.d. (bar) is shown ($n$ = 3; two-way ANOVA with Tukey correction of multiple comparisons).

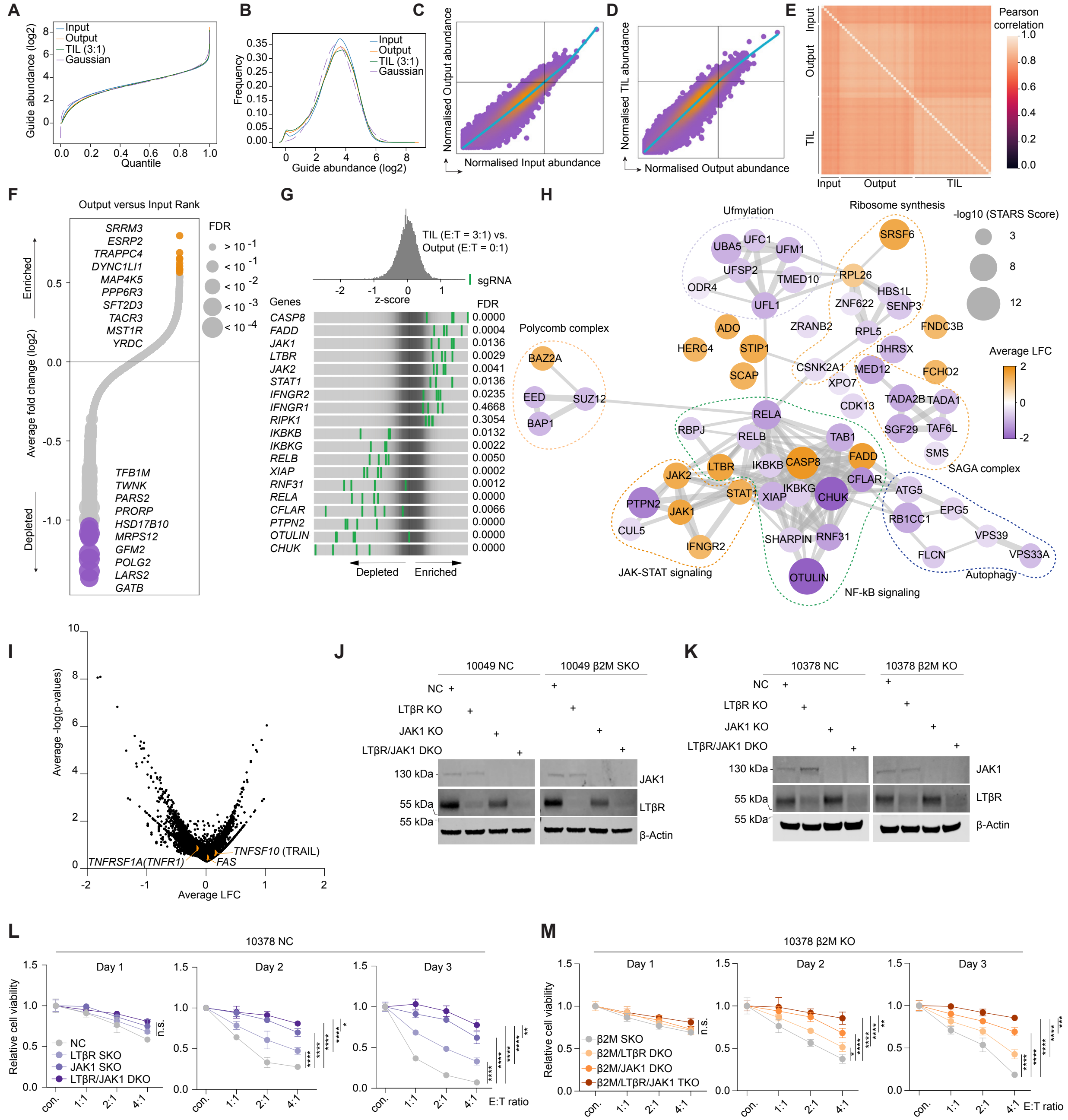

**Supplementary Fig. S4 | LTβR and IFN sensing pathways are involved in TIL-mediated killing. A-B**, Library recovery. Dotted lines represent the Gaussian distribution of (**A**) sgRNA abundance and (**B**) sgRNA frequency (input, output, and TIL-treated groups are shown in solid lines. **C-D**, Effect size model for sgRNAs. A natural cubic spline (solid line) was fit between (**C**) input and output and (**D**) between TIL and output sgRNA abundances, and the effect size is calculated as the residual from the spline. **E**, A matrix of Pearson's correlations of the library distribution from one technical replicate compared to any other technical replicate across input, output, and TIL conditions. **F**, Genes ranked by output versus input-normalized fold change for genome-scale screens, with circle size corresponding to FDR by STARS. Genes with 4-5 mapped guides are visualized; fold changes are calculated only from 4 top-performing guides per gene. Selected top genes with FDR values < = 0.3 at each end are listed and arranged by statistical significance. **G**, Frequency histograms of enrichment or depletion (*z* score) for the selected sgRNAs. sgRNAs targeting indicated genes are shown by the green lines. For genes with more than 4 mapped guides, only the top 4 performing guides were used for fold change and STARS calculations. **H**, STRING network analysis of the depleted and enriched genes with FDR < 0.05, with circles size-scaled by -log$_{10}$(STARS score) and colored by average gene log fold-change. **I**, Volcano plot highlighting the relative depletion and enrichment of sgRNAs targeting *TNFRSF1A (TNFR1)*, *FAS*, and *TNFSF10* (TRAIL). **J-K**, Western blot for JAK1, LTβR, and β-actin in CRISPR-edited **(J)** 10049 and **(K)** 10378 melanoma cells with indicated sgRNA pairs. **L-M**, Viability assessment of CRISPR-edited (**L**) control (sg*NC*) and (**M**) sg*B2M*-null (sg*B2M*) 10378 melanoma cells with indicated sgRNA pairs cocultured with autologous CD8$^+$ TIL at indicated E:T ratios for 1, 2, and 3 days compared to untreated controls. Mean +/- s.d. (bar) is shown ($n$ = 3; Two-way ANOVA with Geisser-Greenhouse correction and Tukey correction for multiple comparisons). *$P$ < 0.05; **$P$ < 0.01; ***$P$ < 0.001; **** $P$ < 0.0001; *ns*, not significant.

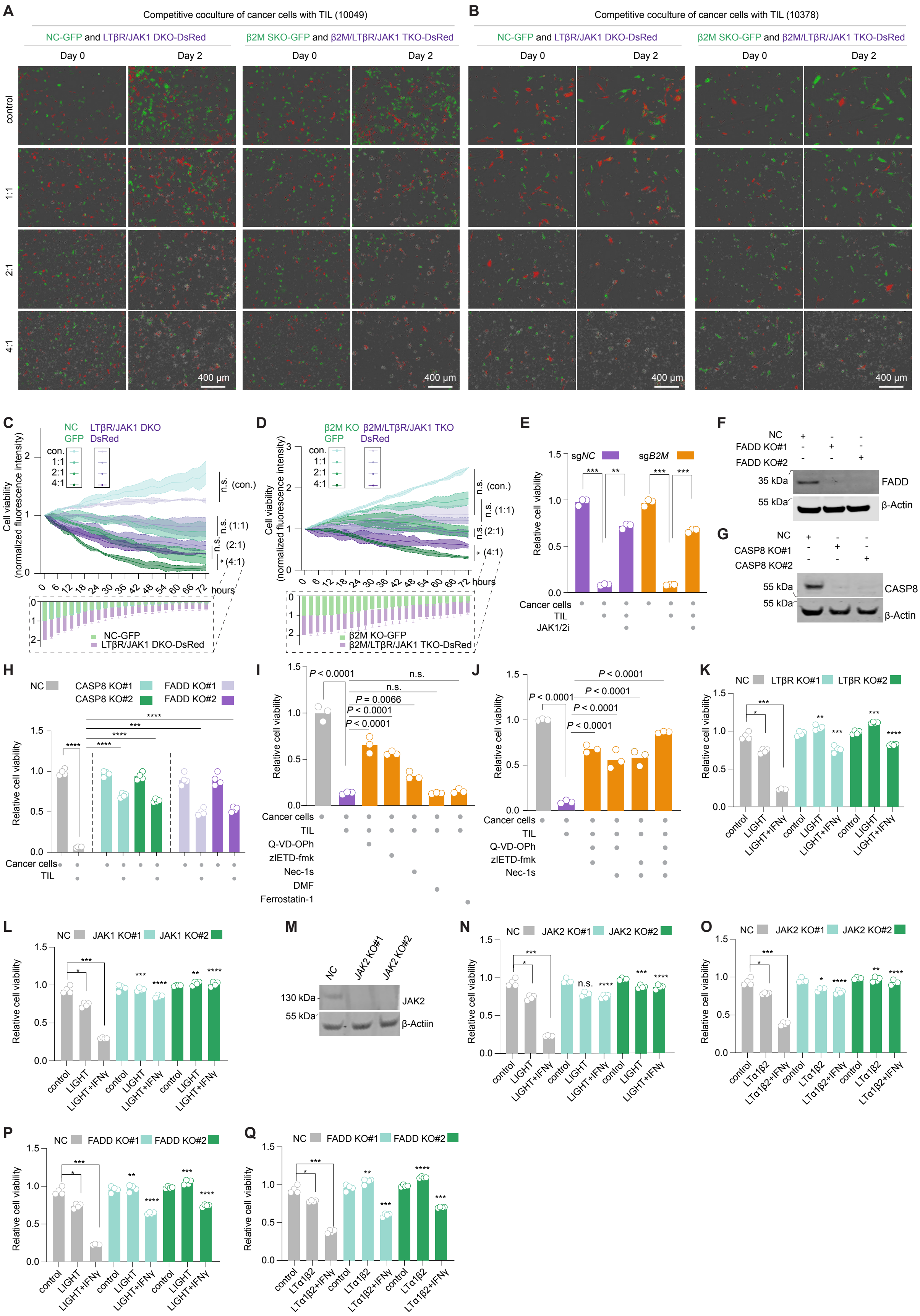


**Supplementary Fig. S5 | See next page for caption.**

**Supplementary Fig. S5 | TIL-mediated killing is via cell-extrinsic apoptosis and necroptosis**. **A,** Representative images from competition assays using equal proportions of (*left*) control (GFP) and LTβR/JAK1 DKO (DsRed) or *B2M*-null (GFP) and (*right*) *B2M*-null/LTβR/JAK1 TKO (DsRed) 10049 melanoma cells co-cultured with autologous TIL at different E:T ratios at indicated times (corresponds to Fig. 3E and 3F). **B,** Representative images from competition assays using equal proportions of (*left*) control (GFP) and LTβR/JAK1 DKO (DsRed) or *B2M*-null (GFP) and (*right*) *B2M*-null/LTβR/JAK1 TKO (DsRed) 10378 melanoma cells co-cultured with autologous TIL at different E:T ratios at indicated times (corresponds to C and D). **C-D**, Competition assays using equal proportions of (**C**) control (GFP) and LTβR/JAK1 DKO (DsRed) or (**D**) *B2M*-null (GFP) and *B2M*-null/LTβR/JAK1 TKO (DsRed) 10378 melanoma cells co-cultured with autologous TIL at different E:T ratios at indicates times. Normalized fluorescence intensity of viable cancer cells (mean +/- s.d.; shaded region) is shown ($n$ = 3, two-way ANOVA with Tukey correction of multiple comparisons). Inset stacked bar blot (E:T = 4:1) depicts proportional decrease in GFP and DsRed cells over time. **E**, Cell viability assessment of control (sg*NC*) and *B2M*-null (sg*B2M*) 10049 melanoma cells pretreated with/without JAK1/2 inhibitor (ruxolitinib, 1 μM) for 3 hours followed by coculturing with autologous TIL (E:T = 4:1) for 2 days compared to untreated control. Individual values (open circles) are shown ($n$ = 3; 2-sided paired $t$-test). **F-G**, Western blot for (**F**) FADD and (**G**) caspase 8 in CRISPR-edited 10049 melanoma cells with 2 different sgRNA per gene target (with β-actin as loading control). **H**, Cell viability assessment of control (sg*NC*), *CASP8*-null (sg*CASP8*), and *FADD*-null (sg*FADD*) 10049 melanoma cells cocultured with autologous TIL (E:T 4:1) for 2 days compared to untreated control. **I-J**, Cell viability assessment of melanoma cells (10049) pretreated (3 hours) with Q-VD-OPh (10 μM), zIETD-fmk (10 μM), Nec-1s (10 μM), DMF (dimethyl fumarate, 25 μM), Ferrostatin-1 (2 μM), or combinations thereof as indicated prior to coculture with autologous TIL (E:T 4:1) for 2 days, compared to vehicle control (0.1% DMSO). Individual values (open circles) are shown ($n$ = 3; one-way ANOVA with Tukey correction of multiple comparisons). **K-L**, Viability assessment of CRISPR-edited 10049 melanoma cells with indicated sgRNA treated with recombinant LIGHT (50 ng/mL), IFNγ (40 ng/mL) alone or in different combinations as indicated for 2 days. **M**, Western blot for JAK2 in CRISPR-edited 10049 melanoma cells with 2 different sgRNA (β-actin as loading control). **N-Q**, Viability assessment of CRISPR-edited 10049 melanoma cells with indicated sgRNA treated with recombinant IFNγ (40 ng/mL) +/- LIGHT (50 ng/mL) or LTα1β2 (50 ng/mL) for 2 days compared to untreated controls. Mean values (bars) and individual values (open circles) are shown ($n$ = 4; 2-sided paired $t$-test). *$P$ < 0.05; **$P$ < 0.01; ***$P$ < 0.001; **** $P$ < 0.0001; *ns*, not significant.

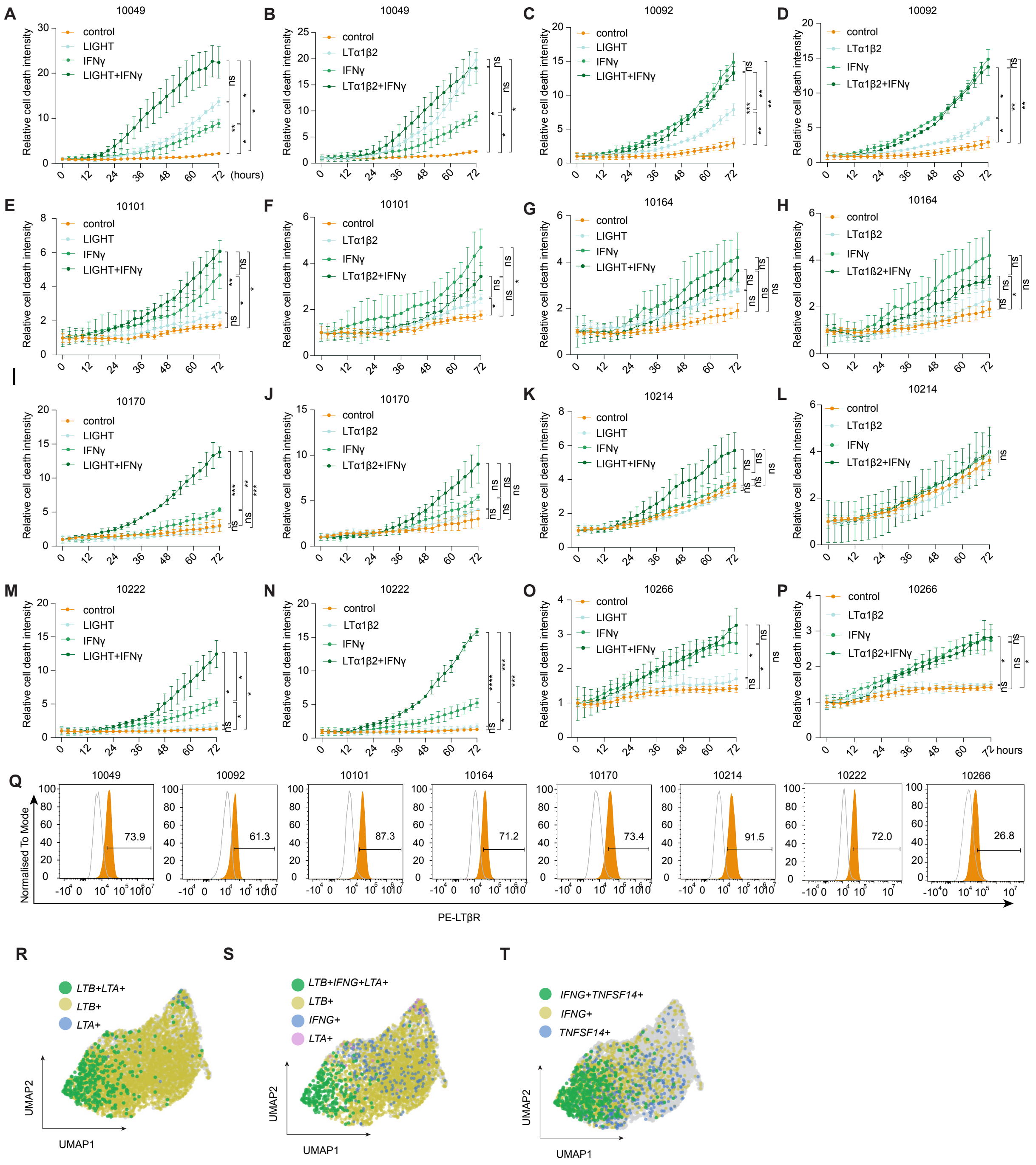

**Supplementary Fig. S6 | Melanoma cell lines show sensitivity to recombinant LTBR ligands and IFNγ**. **A-P**, Cell viability assessment of (**A-B**) 10049, (**C-D**) 10092, (**E-F**) 10101, (**G-H**) 10164, (**I-J**) 10170, (**K-L**) 10214, (**M-N**) 10222, and (**O-P**) 10266 melanoma cell lines treated with recombinant LIGHT (50 ng/mL) or LTα1β2 (50 ng/mL) +/- IFNγ (40 ng/mL) as indicated for 2 days compared to untreated controls. Mean +/- s.d. (bar) is shown ($n$ = 3; Two-way ANOVA with Geisser-Greenhouse correction and Tukey correction for multiple comparisons). **Q**, Flow cytometry histogram of LTβR expression in indicated melanoma cell lines. **R-T**, UMAP embedding depicting the single-cell co-expression of (**R**) *LTB* and *LTA,* (**S**) *LTB*, *IFNG*, and *LTA*, and (**T**) *IFNG* and *TNFSF14* in expanded TIL (10049). *$P < 0.05$; **$P < 0.01$; ***$P < 0.001$; **** $P < 0.0001$; *ns*, not significant.

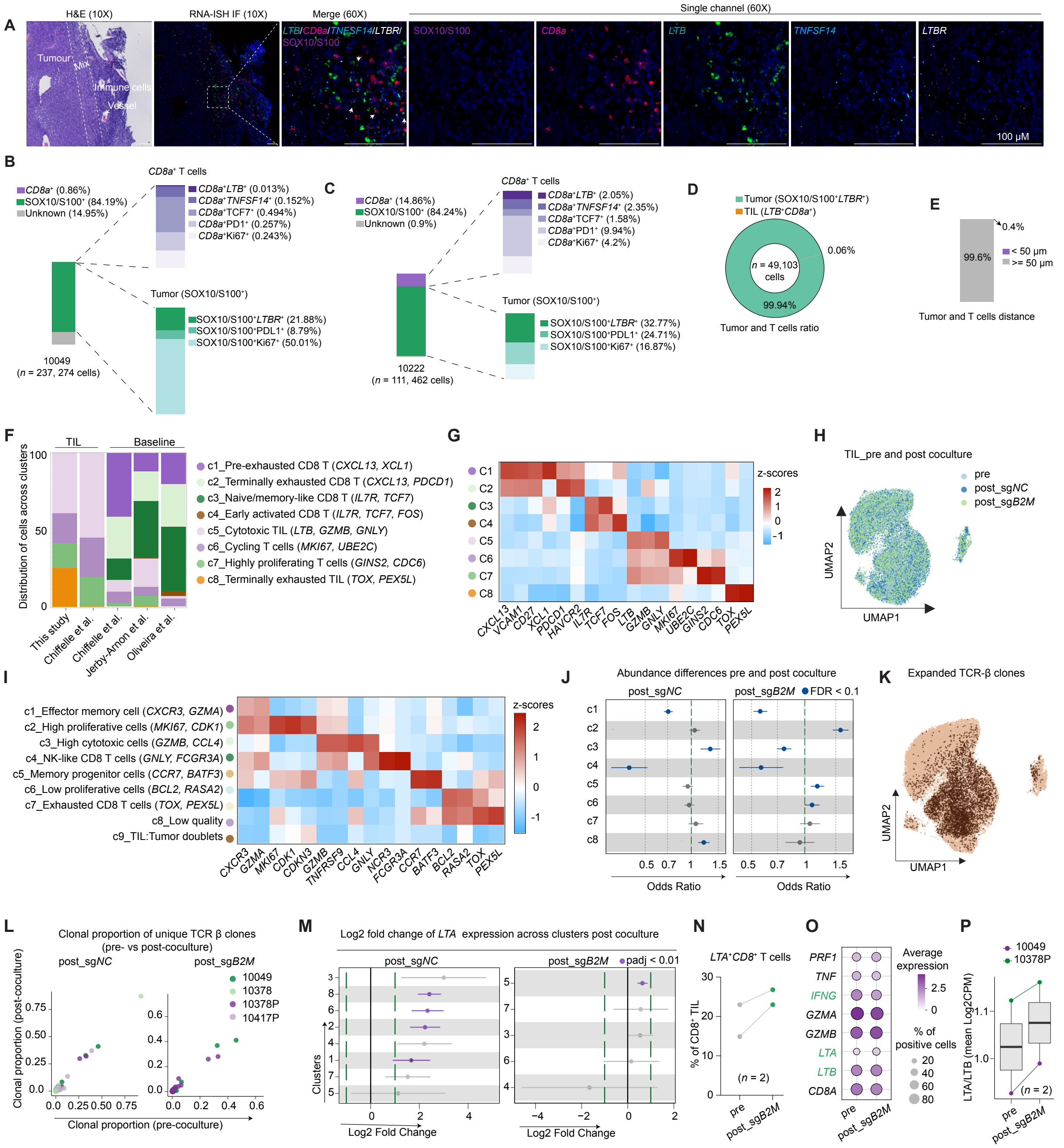


**Supplementary Fig. S7 | Dynamic lymphotoxin expression during expansion and upon coculture with cancer cells**. **A**, H&E and immunofluorescence (IF) with RNA *in situ* hybridization (RNA-ISH) showing the expression of *CD8a*, *LTB*, *TNFSF14*, *LTBR*, and SOX10/S100 in 10049 tumor tissues. **B-C**, Percentage of SOX10/S100$^+$ melanoma cells and *CD8a*$^+$ T cells and their subpopulations in (**B**) 10049 and (**C**) 10222 specimens, respectively. **D**, Percentage of SOX10/S100$^+$*LTBR*$^+$ melanoma cells and *LTB*$^+$*CD8*$^+$ T cells in the cells analyzed in 10049 tumor specimen. **E**, Percentage of SOX10/S100$^+$*LTBR*$^+$ melanoma cells within the indicated distance of *LTB*$^+$*CD8a*$^+$ T cells in the 10049 tumor specimen. **F**, Stacked bar plot showing the relative abundance of cells from various datasets across clusters related to Fig. 4E. **G**, Heatmap showing the representative marker genes differentially expressed across clusters related to Fig. 4E. **H**, UMAP embedding depicting the samples included related to Fig. 5A. **I,** Heatmap showing the representative marker genes differentially expressed across clusters related to Fig. 5A. **J,** Abundance difference in clusters of TIL before and after coculture with sg*NC* (left) and sg*B2M* (right) melanoma cells. **K,** UMAP embedding depicting expanded TCR clones (calculated by TCR β chains). **L**, Scatter plots showing the proportion of TCR clones before and after coculture with sg*NC* (left) and sg*B2M* (right) cancer cells. **M**, Log2 fold changes of *LTA* expression across clusters before and after coculture control (sg*NC*) and *B2M*-null (sg*B2M*) melanoma cells. **N**, Bar plot showing the percentage of *LTA*$^+$*CD8*$^+$ cells in CD8$^+$ TIL before and after coculture with cancer cells transfected with sg*B2M* (*n* = 2). **O**, Dot plot showing selected genes before and after coculture with B2M-nul (sg*B2M*) cancer cells. **P**, Bar plot examining the log2 fold change (mean CPM) of *LTA*/*LTB* ratio in TIL before and after coculture with sg*B2M*.

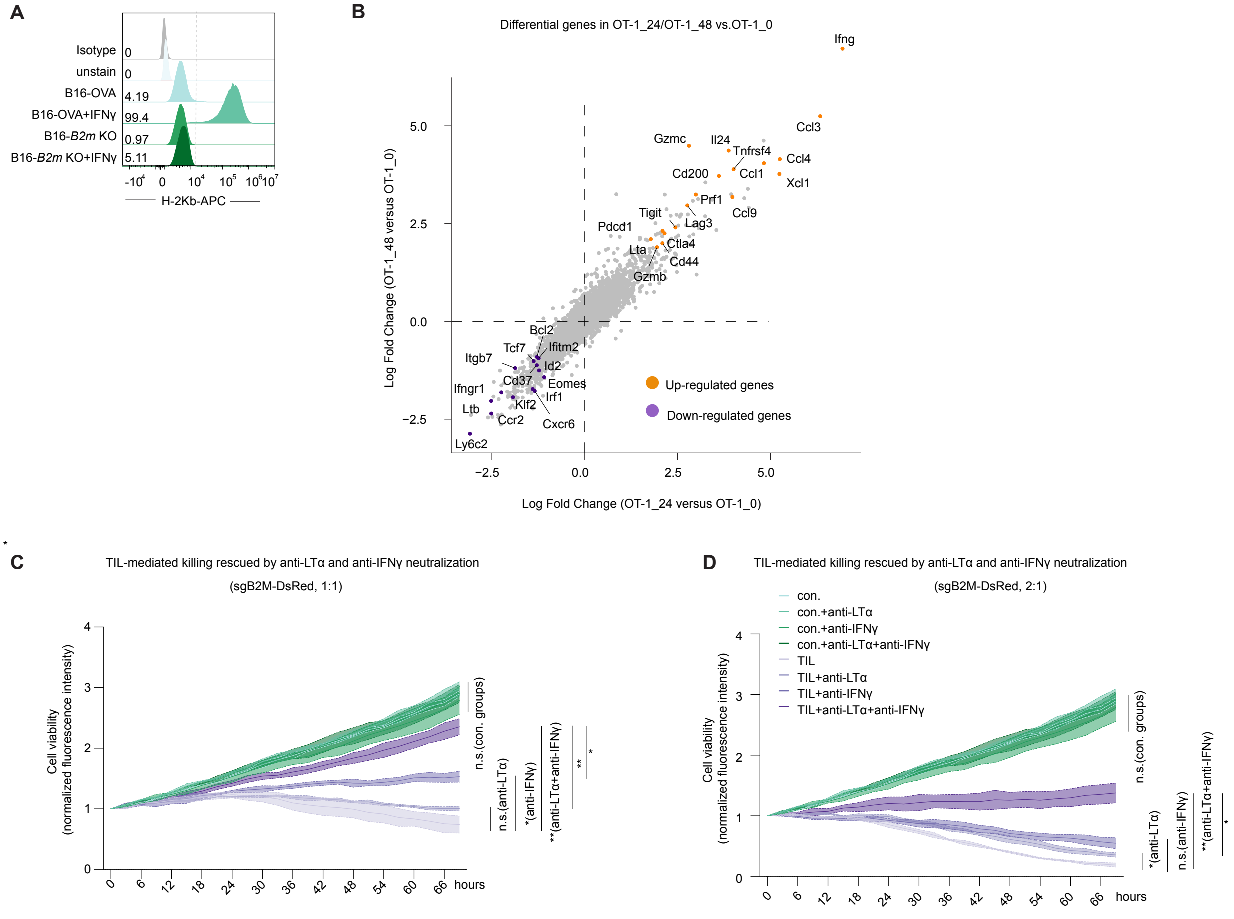


**Supplementary Fig. S8 | Neutralization of LTα and IFNγ rescue cancer cells from TIL-mediated killing.** **A**, Flow cytometry histogram of H-2Kb in B16-OVA or *B2m* KO cells with/without being pretreated with IFNγ (100 ng/mL, 24 hours). **B**, Scatter plot showing the differential gene expression that encodes the surface proteins in OT-1_0 versus OT-1_24 cells (*top*) and OT-1_0 versus OT-1_48 cells (*down*). **C-D,** Normalized fluorescence intensity of viable melanoma cells (10049, sg*B2M*-DsRed) cocultured with autologous TIL and treated with or without the indicated neutralizing antibodies (anti-LTα, 10 µg/mL; anti-IFNγ, 10 µg/mL) at E:T ratio of (**C**) 1:1 or (**D**) 2:1 for the indicated time points.

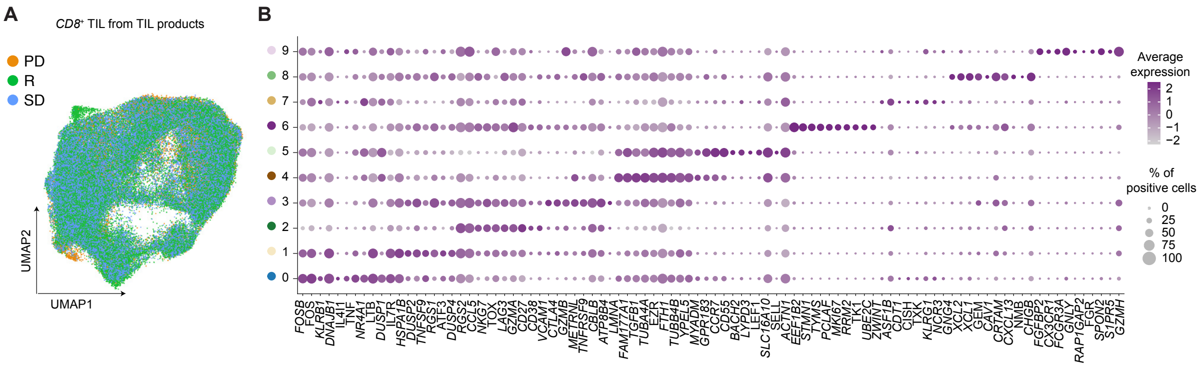


**Supplementary Fig. S9 | The abundance of *LTB*$^{+}$*CD8*$^{+}$ T cells in the expanded TIL products is associated with response to TIL therapy.** **A**, UMAP embedding of scRNA-seq data for TIL depicting the distribution of patients who showed response (R), stable disease (SD), or progression disease (PD) after receiving TIL therapy. **B**, Dot plots showing the representative marker genes differentially expressed across clusters in baseline tumor digests.
